# MicroRNA Profiles in Calcified and Healthy Aorta Differ: Therapeutic Impact of miR-145 and miR-378

**DOI:** 10.1101/736330

**Authors:** Ying Tang, Tapan A. Shah, Edward J. Yurkow, Melissa B. Rogers

## Abstract

Our goal was to elucidate microRNAs (miRNAs) that may repress the excess bone morphogenetic protein (BMP) signaling observed during pathological calcification in the *Klotho* mouse model of kidney disease. We hypothesized that restoring healthy levels of miRNAs that post-transcriptionally repress osteogenic calcific factors may decrease aortic calcification. Our relative abundance profiles of miRNAs in healthy aorta differ greatly from those in calcified mouse aorta. Many of these miRNAs are predicted to regulate proteins involved in BMP signaling and may control osteogenesis. Two differentially regulated miRNAs, miR-145 and miR-378, were selected based on three criteria: reduced levels in calcified aorta, the ability to target more than one protein in the BMP signaling pathway, and conservation of targeted sequences between humans and mice. Forced expression using a lentiviral vector demonstrated that restoring normal levels repressed the synthesis of BMP2 and other pro-osteogenic proteins and inhibited pathological aortic calcification in *Klotho* mice with renal insufficiency. This study identified miRNAs that may impact BMP signaling in both sexes and demonstrated the efficacy of selected miRNAs in reducing aortic calcification *in vivo*. Calcification of the aorta and the aortic valve resulting from abnormal osteogenesis is common in those with kidney disease, diabetes, and high cholesterol. Such vascular osteogenesis is a clinically significant feature. The calcification modulating miRNAs described here are candidates for biomarkers and “miRNA replacement therapies” in the context of chronic kidney disease and other pro-calcific conditions.

## Introduction

The American Heart Association projects that 45% of Americans will have some form of heart and/or vascular disease by 2035 (5). Calcification of blood vessels reduces their diameter and plasticity and ultimately promotes ischemic events (22). Risk factors that promote both vascular and valvular calcification include aging, renal failure, and male sex; as well as hypercholesterolemia, smoking, and diabetes (46, 54). An aging population and the epidemic of obesity ensure that cardiovascular disorders will be a major public health threat for decades to come. Understanding the signaling processes that regulate the behavior of vascular cells will reveal novel approaches to preventing and treating these devastating pathologies. Here we focused on identifying microRNAs (miRNAs) that regulate pathological cardiovascular calcification.

The BMP signaling pathway is essential for osteogenesis and vascular calcification (8, 16, 46). BMP ligands bind BMP receptors (BMPR), which phosphorylate and activate SMAD signaling. SMAD signaling increases transcription of *Runx2* and *Msx2*, which are master osteogenic transcription factors that promote osteogenic differentiation (16, 25, 46). Abnormally elevated levels of the pro-osteogenic bone morphogenetic protein 2 (BMP2) and increased BMP signaling are implicated in all forms of pathological cardiovascular calcification. For example, BMP2 is synthesized in human atherosclerotic plaques (7) and calcified stenotic valves (33). Moreover, BMP2 can induce the pro-osteogenic RUNX2 and MSX2 proteins in valve and aortic cells and can induce calcification and ossification both *in vivo* and *in vitro* (7, 10, 30, 32, 34, 36, 37, 49, 51, 53). Genetic deficiency and pharmacological inactivation of BMP signaling reduced calcification not only in cultured cells (4, 17, 49) but also in several mouse models including *Klotho* mutant mice with renal failure (14, 17, 28).

Given the importance of BMP signaling in cardiovascular calcification, identifying post-transcriptional regulators of BMP signaling may provide new insights and targets for the treatment of cardiovascular disease. To increase potential clinical translatability, we also considered evolutionary conservation. The 3’-untranslated region (3’-UTR) of messenger RNAs (mRNAs) binds regulatory proteins and miRNAs that influence polyadenylation, translation efficiency, and stability of mRNAs (12). All BMP ligands may be post-transcriptionally regulated. However, the extreme evolutionary conservation of the *Bmp2* 3’-UTR relative to other BMP ligand 3’-UTRs supports a potentially greater role for post-transcriptional *Bmp2* regulatory mechanisms that are conserved between mouse and man (15, 39, 43). MicroRNAs are crucial regulators in cardiovascular pathologies (21). The goal of this study was to compare the profiles of miRNAs that modulate the synthesis of BMP2 and downstream BMP signaling proteins in the aorta of healthy mice relative to mice with pathologically calcified aorta. Our mouse model is a *Klotho* hypomorph that causes dysregulated mineral metabolism (24).

The phenotype of mice homozygous for *Klotho* mutations includes a short lifespan, atherosclerosis, ectopic calcification, and renal disease. The KLOTHO protein, along with 1,25(OH)_2_ vitamin D3 and FGF23, tightly regulates phosphate homeostasis (19, 50). Deficiency of KLOTHO leads to renal disease and subsequent hyperphosphatemia and ectopic calcification of soft tissues such as aorta, aortic valves and kidneys (17, 50). Restoration of normal KLOTHO levels ameliorates calcification (19). The dramatic calcification of the aorta and other tissues in *Klotho* mutant mice within 6-7 weeks of birth make them experimentally attractive for studying gene regulation and signaling changes during pathological calcification. Understanding the mechanism of aortic calcification in the context of kidney failure is important because 17% of the population over the age of 30 may suffer from this major cardiovascular disease risk factor by 2030 (5).

Our objective was to elucidate a comprehensive profile of miRNAs whose abundance is altered in the calcified aorta of *Klotho* homozygous mutant mice with renal disease. Our miRNA profiles from both male and female healthy and diseased mice reflect the extensive changes that occur during vascular disease. Here we discuss the subset of differentially regulated miRNAs that may impact the pathological osteogenesis that contributes to vascular calcification. We also demonstrate that selected miRNAs can modulate the BMP2 ligand and proteins involved in BMP signaling and calcification *in vivo*. Finally, we provide proof-of-principle evidence that increasing the abundance of miRNAs that inhibit BMP signaling can ameliorate aortic calcification. This study begins to fill a key gap in our understanding of post-transcriptional processes that control BMP signaling and calcification in vascular tissues and provides experimental support for miRNA replacement therapies in cardiovascular disease.

## Material and Methods

### Tissue collection

Mice bearing the *Klotho* mutation were a gracious gift from Dr. Makoto Kuro-o (Jichi Medical University) by way of Dr. Sylvia Christakos (Rutgers New Jersey Medical School). The mice were a mixture of strains FVB, C57Bl/6J, and C3H/J. Because homozygous *Klotho* null mice are infertile, mice were maintained by heterozygous-by-heterozygous mating. 25% of each litter was homozygous and exhibited the full *Klotho* phenotype, including ectopic soft tissue calcification. 50% were heterozygous and 25% were wild type. No statistically significant differences in any parameter were observed between mice bearing the heterozygous and wild type *Klotho* genotypes (Table S1). Assessed parameters included body and organ weights, gene expression, and calcium levels. Because our analyses were consistent with published data indicating that the *Klotho* mutation is fully recessive, heterozygous and wild type samples were presented together as “healthy control” samples. In all experiments, tissues from both male and female mice were assayed. The resulting data were presented with closed and open circles indicating male and female values, respectively.

All animal procedures were in accordance with the guidelines for Care and Use of Experimental Animals and approved by the NJ Medical School Institutional Animal Care and Use Committee (IACUC protocol #PROTO999900898). Control and *Klotho* homozygote mice were fed regular chow and euthanized at 50 + 1 days of age. After weaning, *Klotho* homozygotes received softened chow on the floor of the cage. On the day of necropsy, mice were killed with an inhalation overdose of isoflurane. Immediately thereafter, the heart was perfused *via* the left ventricle with phosphate buffered saline (PBS, pH 7.3), to remove excess blood. The aorta, including the ascending aorta, arch and descending thoracic aorta, was removed. After cutting at the surface of the heart, the aorta was rinsed in PBS, blot dried and weighed. The aorta was snap-frozen in liquid nitrogen and stored at −80◦C. Frozen tissues were ground in liquid nitrogen using a mortar and pestle. To facilitate handling small tissues such as the diseased aortas from *Klotho* homozygotes, acid-washed glass beads (Millipore-SIGMA, St. Louis, MO, # G1277) were added during the grinding to both mutant and control tissues. Glass beads did not affect the biochemical assays (Fig. S1A, B).

### Western Blots

Frozen ground tissue was solubilized in RIPA buffer, sonicated, and subjected to western blot analyses as described in Shah *et al.* (43). BMP signaling was measured using a monoclonal phospho-SMAD 1/5/9(8) antibody (Cell Signaling Technology, Danvers, MA, #13820) at a dilution of 1:1000. The pSMAD antibody was authenticated as described in Fig. S1C and D. Polyclonal total SMAD 1/5/9(8) (Santa Cruz Biotechnology, Inc., Santa Cruz, CA, #sc-6031-R) and a polyclonal actin antibody (Santa Cruz Biotechnology, Inc., Santa Cruz, CA, #sc-1615-R) were used at a dilution of 1:1000. Both pSMAD and tSMAD antibodies recognize the three BMP specific SMADs 1, 5, 9. In all cases, the secondary antibody was Goat Anti-Rabbit HRP (Abcam, Cambridge, MA, # ab97080) at a dilution of 1: 20,000. Antibody-bound proteins were detected using SuperSignal™ West Femto Maximum Sensitivity Substrate (ThermoFisher Scientific, Waltham, MA, # 34096) and imaged using a FluoroChem M (Protein Simple, San Jose, California).

### Calcium assays

Ground tissue was solubilized and lysed by sonication on ice in PBS, pH 7.3 containing 0.16 mg/mL heparin. The Cayman Chemical Calcium Assay kit was used to measure calcium levels (Ann-Arbor, MI, #701220). Calcium levels were normalized to protein levels measured using the Bradford assay (Bio-Rad Laboratories, Hercules, CA, # 5000006).

### Spatial mapping of calcified structures

The patterns of mineralization in *Klotho* heterozygote *vs. Klotho* mutant mice were determined using microcomputerized tomography (microCT) at the Rutgers Molecular Imaging Center (http://imaging.rutgers.edu/). Mice were scanned using the Albira® PET/CT (Carestream, Rochester, NY) at standard voltage and current settings (45kV and 400μA) with a minimal voxel size of <35 μm. Voxel intensities in the reconstructed images were evaluated and segmented with VivoQuant image analysis software (version 1.23, inviCRO LLC, Boston).

### MicroRNA microarray and data analysis

Aortic samples were obtained from both male and female control and *Klotho* mutant mice at 7 to 8 weeks of age (Table 1). RNA was extracted using the miRNeasy Mini Kit (Qiagen Inc., Germantown, MD, # 217004). The RNA quality was checked using a Bioanalyzer (Agilent Technologies, Santa Clara, CA).

**Table 1.** Number of mice (n) used for the miRNA profiling

|  | **Males** | **Females** |
| --- | --- | --- |
| **Group** | **n** | **n** |
| Control (Wt) | 5 | 4 |
| Control (K+) | 2 | 1 |
| *Klotho* mutant (KK) | 5 | 2 |

MicroRNA expression in the aorta was assessed with the Applied Biosystems™ GeneChip™ miRNA 4.0 Array (ThermoFisher Scientific, Waltham, MA, #902412) in the Rutgers NJMS Genomics Center. The GeneChip™ miRNA 4.0 Array contained the miRNAs and pre-miRNAs (precursors to the miRNAs) listed in the Sanger miRBase v20. The miRNA arrays were processed following the manufacturer’s instructions.

Briefly, using the Applied Biosystems™ FlashTag™ Biotin HSR RNA Labeling Kit (ThermoFisher Scientific, Waltham, MA, the # 902446), 400 ng of total RNA was labeled with biotin and hybridized to the miRNA 4.0 Array (ThermoFisher Scientific, Waltham, MA, #902412) for 18 hours at 48°C using an Affymetrix® 450 Hybridization Oven (Affymetrix, Santa Clara, CA). After washing and staining on an Affymetrix® 450 Fluidics Station (Affymetrix, Santa Clara, CA), using the Applied Biosystems™ GeneChip™ Hybridization, Wash, and Stain Kit (ThermoFisher Scientific, Waltham, MA, #900720), the arrays were scanned using the Affymetrix® Scanner 3000 7G (Affymetrix, Santa Clara, CA). CEL files were generated using the Affymetrix data extraction protocol in the Affymetrix GeneChip® Command Console® Software (Affymetrix, Santa Clara, CA, USA). The resulting CEL files (GSE135759) were analyzed using the Partek Genomics Suite 7.0 (Partek Inc., St. Louis, MO), including the Robust-Multi-array Average algorithm to correct for microarray background and to normalize miRNA expression profiles on a log scale. Partek Genomic’s ANOVA analysis identified the miRNAs whose abundance differed significantly between control and *Klotho* homozygous mutant mice. Minimal significance was defined as a *p* value of 0.05.

### Bioinformatic analyses

MicroRNAs that were significantly down-regulated in both sexes of *Klotho* homozygous mice compared to controls were (further) used to perform target prediction (see microarray data analysis). Predicted target profiles were obtained from the DNA Intelligent Analysis (DIANA)-micro T-CDS (v. 5.0) which permits the entry of multiple miRNAs simultaneously (http://www.microrna.gr/webServer) (38). The micro T-CDS threshold score for predicted targets was set at 0.6. Scores ranged from 0 to 1, with a higher score indicating an increased probability of a true microRNA target.

Alignments between selected miRNAs and mouse genes were predicted by MiRanda (6) and TargetScan 7.2 (http://www.targetscan.org/mmu_72/ (1)).

### Real-Time Reverse Transcription PCR for detection of microRNA-145 and microRNA-378a and target gene expression

MicroRNA cDNA was synthesized using the miScript PCR Kit (Qiagen Inc., Germantown, MD, # 218073). The miScript PCR kit utilized a universal oligo-dT primer tag to convert all RNA species (total RNA and miRNA) into cDNA. A universal reverse primer supplied with the miScript PCR kit was used to amplify cDNA from each miRNA. The miRNA forward primers were designed based on miRNA sequences in miRBase 22 (http://www.mirbase.org/). GCG or CGC was added to the 5’ end of the primers when the GC % was low. NCBI nucleotide blast was used to confirm the absence of non-specific binding sites for the forward primers. Messenger RNA (mRNA) cDNA was synthesized using the QuantiTect® Reverse Transcription kit (Qiagen Inc., Germantown, MD, # 205313). Intron-spanning primers for mRNA were used to eliminate amplicons generated from any contaminating genomic DNA. Primer sequences are shown in Table 2.

**Table 2.**
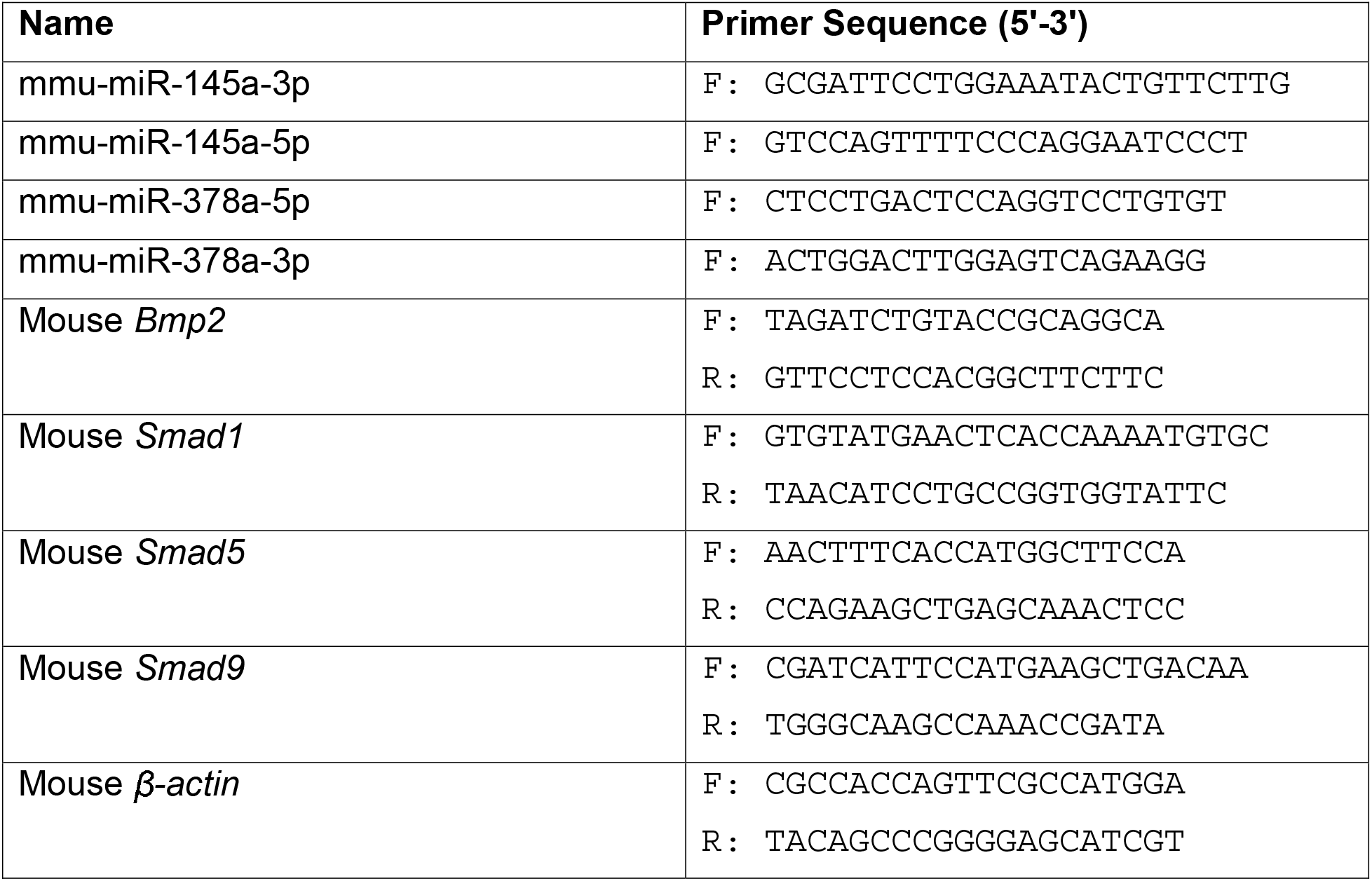
Reverse Transcription PCR primer sequences

Quantitative PCR was performed on a CFX96 Touch™ Real-Time PCR Detection System (Bio-Rad Laboratories, Hercules, CA, #1855196) using the QuantiTect® SYBR® Green PCR kit (Qiagen Inc., Germantown, MD, #204145) under the following conditions: 15 min at 95°C, 39 cycles of 15 seconds at 94°C, 30 seconds at 55°C and 30 seconds 70°C. MicroRNA abundance was normalized to U6 and mRNA expression was normalized to actin. The human U6 primer used in this experiment was included in the miScript Primer assay kit (Qiagen Inc., Germantown, MD, #218300).

### Lentivirus generation and titering performed by Biosettia, Inc. (San Diego, CA)

Mouse miRNA mimics (pLV-Sm22a-[mmu-mir-145], 1.7×10^7^ IU/ml; pLV-Sm22a-[hsa-mir-378a], 2.1×10^7^ IU/ml) or control (pLV-Sm22a-[mir-ctrl], 1.7×10^7^ IU/ml) bearing 3^rd^ generation vectors (Vesicular Stomatitis Virus (VSV) G pseudotype) were purchased from Biosettia, Inc. The mature mir-378a sequences generated from the human precursor are identical to the mouse mature sequences. General information regarding Biosettia self-inactivated lentiviral vectors can be found at https://biosettia.com/mirna/. Briefly, the miRNA precursors were cloned within the intron of the human EF1a promoter region in which the EF1a promotor was replaced by the mSm22a promoter. The pLV-microRNA vectors were cotransfected with three helper plasmids (Gag-pol, Rev and VSV-G) and measured using the antibiotics selection method (blasticidin). Synthesis and titering are described at https://biosettia.com/download/protocols/biosettia-mirna-lentiviral-expression-vector-manual.pdf.

### Lentiviral microRNA-145 and microRNA-378a transduction

Lentivirus vectors expressed the mir-145 or mir-378a precursor RNAs under the control of the vascular smooth muscle cell (VSMC)-specific mSm22a promoter. This well-characterized minimal promoter directs gene expression specifically in VSMCs of large and medium-sized arteries in adult mice, with no expression in other muscle cells or organs including the kidney and liver (11, 26, 27, 31). The negative control was the empty virus vector. Newly weaned *Klotho* mutant homozygotes were injected with empty lentivirus or lentivirus bearing mir-145 or mir-378a *via* their tail veins on 5 alternate days. The total lentivirus achieved in each mouse at the end of the treatment window was 4×10^5^ IU/g body weight. The mir-145 lentivirus generates two mature miRNAs: miR-145a-5p and miR-145a-3p. The mir-378a lentivirus generates two mature miRNAs: miR-378a-5p and miR-378a-3p.

### Statistical Analysis

Data were analyzed with GraphPad (version 7; GraphPad Software Inc, La Jolla, CA) and are presented as mean ± Standard Error Measurement (SEM) as indicated. Statistical significance was assessed with the Student’s two-tailed t-test. Differences were considered significant if *p*<0.05. The significance of any differences in male and female values was assessed by a t-test. With one exception, the male and female RT-PCR, western blot, and calcium values did not significantly differ. Consequently, male and female values are indicated as closed or open circles and were pooled for statistical calculations. The one exception, miR-378a-3p levels, are noted in the results and figure legends.

## Results

### BMP signaling and calcification is elevated in aorta from Klotho homozygous mutant mice

BMP signaling is required for valve calcification in *Klotho* null mice (17). We assessed BMP signaling in the calcified aorta of mice homozygous for the original hypomorphic *Klotho* allele (24). BMP signaling and calcium levels in aorta from control mice that were either wild type (*Kl*^*+/+*^) or heterozygous (*Kl*^*kl*/*+*^) for the *Klotho* mutation were compared to the aorta from *Klotho* mutant homozygotes (*Kl^kl/kl^*). BMP signaling, as assessed by phosphorylation of the three BMP-specific SMADs 1, 5, and 9, was induced nearly 2-fold in aorta from male and female *Klotho* homozygous mice relative to control mice (Fig. 1A, B; *p* = 0.002). Calcium levels in the aorta from *Klotho* mutant homozygotes also were elevated by over 2-fold relative to control aorta (Fig. 1C, p<0.0001). PET-CT imaging revealed profound mineralization in the aortic sinus and ascending aorta of *Klotho* mutant (*Kl^kl/kl^*) mice, but not in a control heterozygous (*Kl*^*kl*/*+*^) littermate (Fig. 1D, E). Together, these results confirmed that homozygosity for the hypomorphic *Klotho* mutation amplifies BMP signaling and calcification in the aorta.

**Fig. 1.**
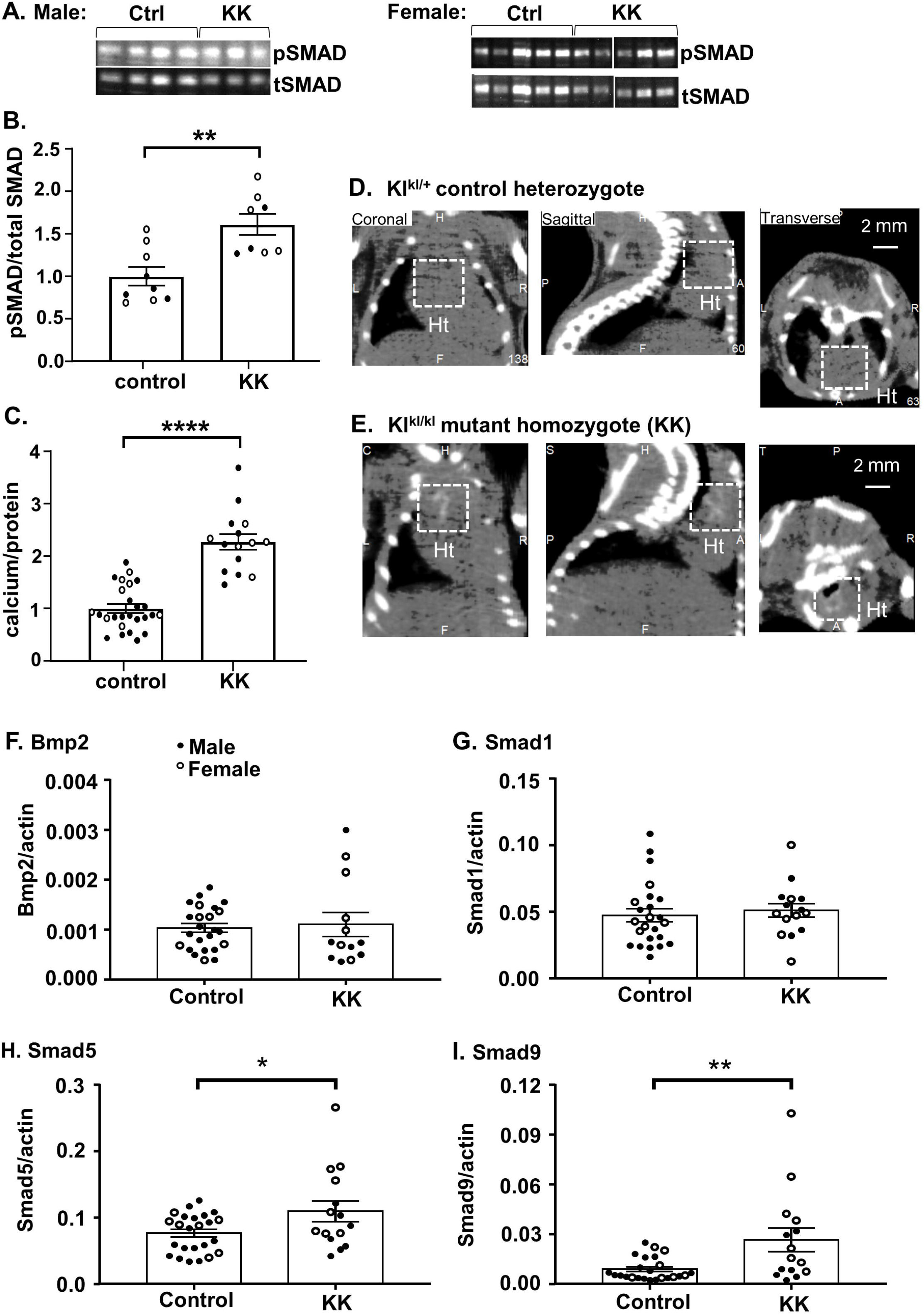
BMP signaling and calcification levels in aorta from homozygous *Klotho* mutant mice. Aortas were isolated from control (Ctrl) mice with normal kidney function (*Kl*^*+/+*^ and *Kl*^*kl*/*+*^) or mice homozygous (*Kl^kl/kl^*) for the *Klotho* mutation with renal disease (KK). Average parameter values are presented with standard error measurements (SEM). **A.** BMP signaling levels were assessed using an antibody that detects the phosphorylated forms of SMADs 1, 5, and 9 (pSMAD). Representative western blot panels showing pSMAD1/5/9 and total SMAD1/5/9 levels, males (left) and females (right). All female samples were loaded on one gel; however, one mis-loaded lane was excised from the image. **B.** pSMAD1/5/9 levels were normalized to total SMAD1/5/9 (tSMAD). Solid and hollow circles represent individual male and female values, respectively. **C.** Average calcium levels normalized to protein levels. **D, E.** Male littermates *w*ere scanned with an Albira PET/CT Imaging System (Carestream, Rochester, NY) set at 45 kV, 400 µA, and <35 μm voxel size. Voxel intensities in the reconstructed images were evaluated and segmented with VivoQuant image analysis software (version 1.23, inviCRO LLC, Boston MA). The dashed white lines mark mineralized areas of the aortic sinus and ascending aorta present in the heart (**Ht**) of the *Klotho* mutant homozygote **(E)**, but not in the heterozygous control *Kl*^*kl*/*+*^ **(D)**. All images are shown at the same scale. Selected RNAs were measured by RT PCR and normalized to actin RNA levels. *Bmp2* **(F)** and *Smad1* **(G)** RNA levels did not change. *Smad5* **(H)** and *9* **(I)** RNA levels were significantly increased relative to control mice. In all panels, the bars indicate the mean value +/− SEM. * *p* < 0.05, ** *p* < 0.01, **** *p* < 0.001. Values did not differ significantly between sexes. All experiments were repeated at least twice with similar results.

We then tested the hypothesis that increased abundances of aortic RNAs encoding the ligand BMP2 or the BMP signaling intermediaries SMADs 1, 5, or 9 accounted for the increased signaling in the *Klotho* homozygotes. The levels of the *Bmp2* and *Smad1* RNAs were similar to those in control mice (Fig. 1F, G). In contrast, *Smad5* and *9* RNA levels were 1.4-fold and 3.0-fold higher, respectively, in the aorta of *Klotho* homozygotes compared to control mice (Fig. 1H, I, p < 0.05 and p < 0.01). Although we have yet to rule out differential translation of the *Bmp2* and *Smad1* RNAs, we did not observe disease-associated differences in mRNA abundances. However, the increased abundance of the RNAs encoding the SMAD5 and 9 intracellular BMP signaling intermediaries indicates that increased transcription and/or stability of messages that encode BMP signaling proteins occurs in the aorta of *Klotho* mutant homozygotes.

### MicroRNA profiles differ in the aorta from Klotho homozygous mutant mice

BMP signaling and calcium levels are doubled in the aorta of Klotho homozygotes (Fig. 1). Alterations in miRNAs that target pro-calcific ligands such as BMP2 or signal intermediaries such as the SMADs may alter translational efficiency or destabilize mRNAs leading to changes in abundance. Consequently, we hypothesized that miRNA profiles differ in aorta from healthy control mice and *Klotho* homozygous mutant mice with dysregulated mineral metabolism. To identify miRNAs involved in aortic calcification, we profiled miRNAs in aortas from control healthy and *Klotho* mutant homozygotes. In total, 145 miRNAs were significantly increased, and 106 miRNAs were significantly decreased in *Klotho* mutant homozygous males relative to control mice (p<0.05, Fig. 2A, Table 3, Table S2, GSE135759). In *Klotho* mutant females, 121 miRNAs were significantly increased, and 92 miRNAs were significantly decreased (p<0.05, Fig. 2B, Table 3, Table S2, GSE135759). The abundances of 31 miRNAs were altered significantly in both male and female *Klotho* homozygous mutant mice relative to control mice (p<0.05, Fig. 2C, Table 4). Of these 10 miRNAs were up-regulated and 12 miRNAs were down-regulated in both sexes of *Klotho* homozygous mice. These results are consistent with a major role for miRNA-mediated post-transcriptional regulation in aortic calcification.

**Table 3.** The number and percentage of miRNAs that differed in control relative to *Klotho* homozygotes (KK), p<0.05

|  | **Male** | **Female** |
| --- | --- | --- |
| **Any significant change** |  |  |
| Increased in KK | 145 (7.6%) | 121 (6.3%) |
| Reduced in KK | 106 (5.6%) | 92 (4.8%) |
| **1.5-fold difference** |  |  |
| Increased in KK | 97 (5.1%) | 71 (3.7%) |
| Reduced in KK | 60 (3.1%) | 50 (2.6%) |
| **2-fold difference** |  |  |
| Increased in KK | 77 (4.0%) | 48 (2.5%) |
| Reduced in KK | 25 (1.3%) | 28 (1.5%) |

**Table 4.**
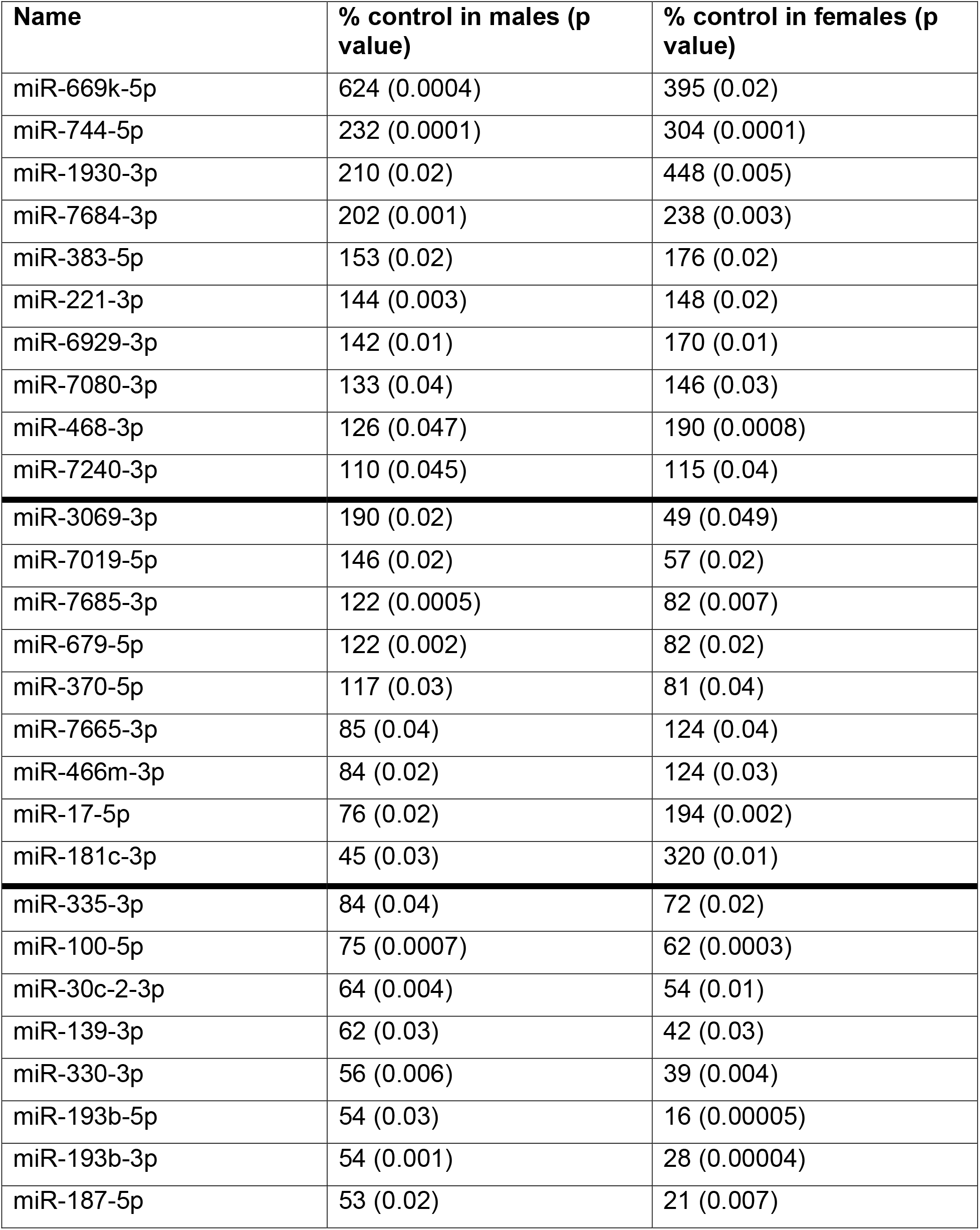
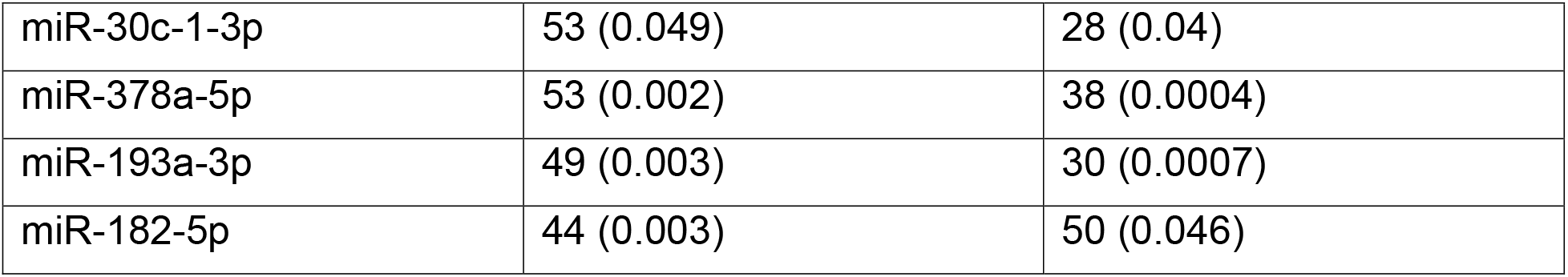
Relative abundance of miRNAs that were differentially expressed in both male and female *Klotho* homozygous mutant mice as assessed by microarray comparisons.

**Fig. 2.**
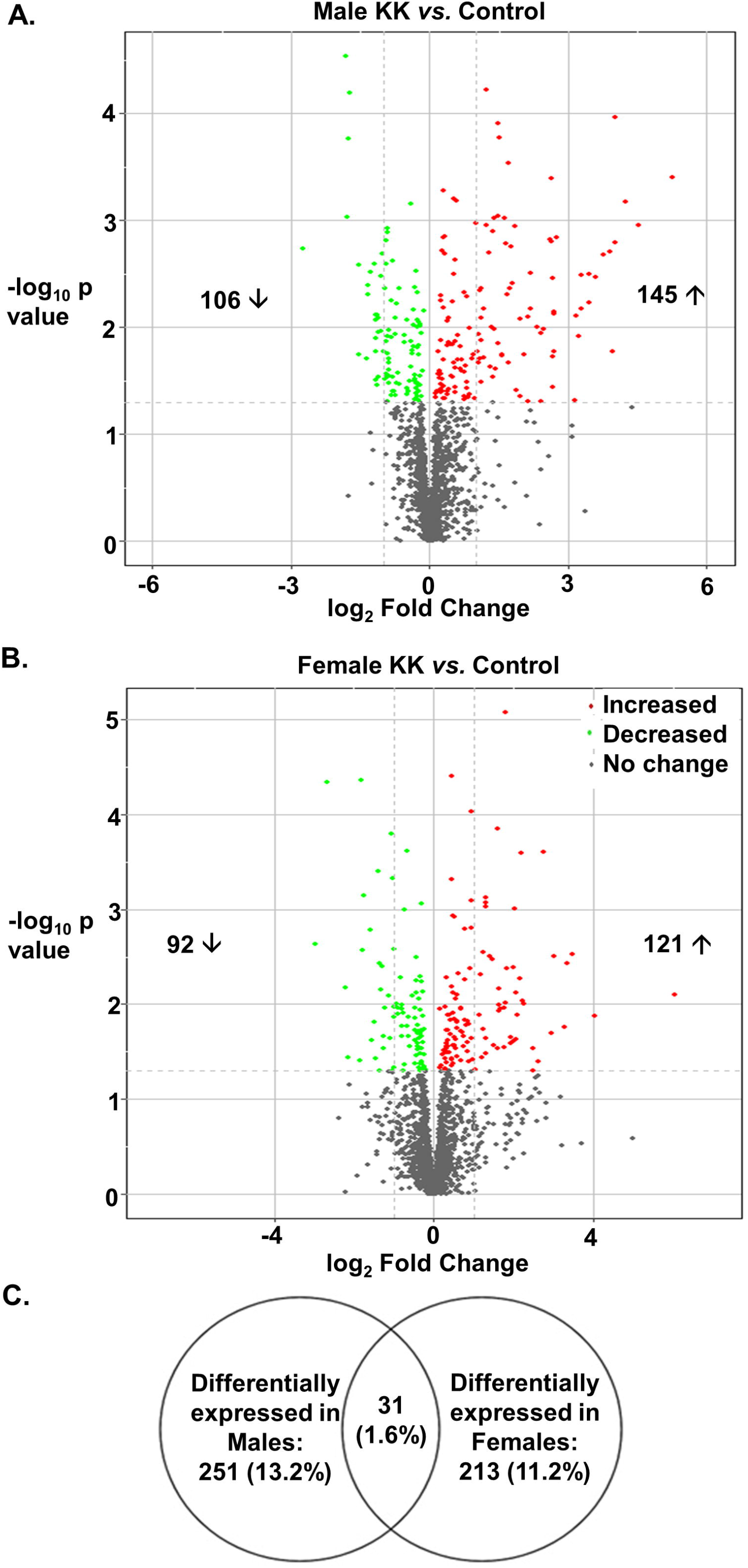
MicroRNA profiles in the aorta from control and *Klotho* homozygous *(KK)* mutant mice. RNA was extracted from the aorta of healthy controls (*Kl*^*+/+*^ and *Kl*^*kl*/*+*^) or *Klotho* homozygous (*Kl^kl/kl^*) mice (KK). Samples were assayed using Affymetrix miRNA version 4.0. Data analysis was performed after Robust Multi-Array Average (RMA) normalization with Partek Genomics Suite 7.0. **A, B.** Volcano plots illustrate miRNAs whose abundances differed significantly in aorta from diseased mice compared to control mice (*p* < 0.05). The X axis indicates log_2_ fold change and the Y axis indicates - log_10_ p value. Vertical dotted lines indicate 2-fold change. **C.** The Venn diagram shows the number and percentage of miRNAs differentially expressed in male or female mice or in both sexes from a total of 1908 miRNAs.

Curiously, the abundances of 9 miRNAs were altered by *Klotho* homozygosity but in the opposite directions in each sex: miR-3069-3p, miR-7019-5p, miR-7685-3p, miR-679-5p and miR-370-5p were up-regulated in males and down-regulated in females, whereas miR-7665-3p, miR-466m-3p, miR-17-5p and miR-181c-3p were down-regulated in males and up-regulated in females (Table 4). The focus of this study was to restore natural repressive mechanisms whose weakening may contribute to pathological calcification. Therefore, we focused on miRNAs whose abundance is reduced in the calcified aorta of both sexes.

A manually compiled list of BMP signaling-relevant genes predicted by TargetScan to be targeted by selected miRNAs is shown in Table 5. Together, these results suggest the disease-associated miRNA profile in *Klotho* homozygotes may influence the tendency of aortic cells to undergo osteogenic differentiation. We postulate that microRNAs that target more than one member of the BMP signaling pathway may inhibit BMP signaling by coordinately down-regulating multiple proteins and effectively restrain pathological calcification.

**Table 5.**
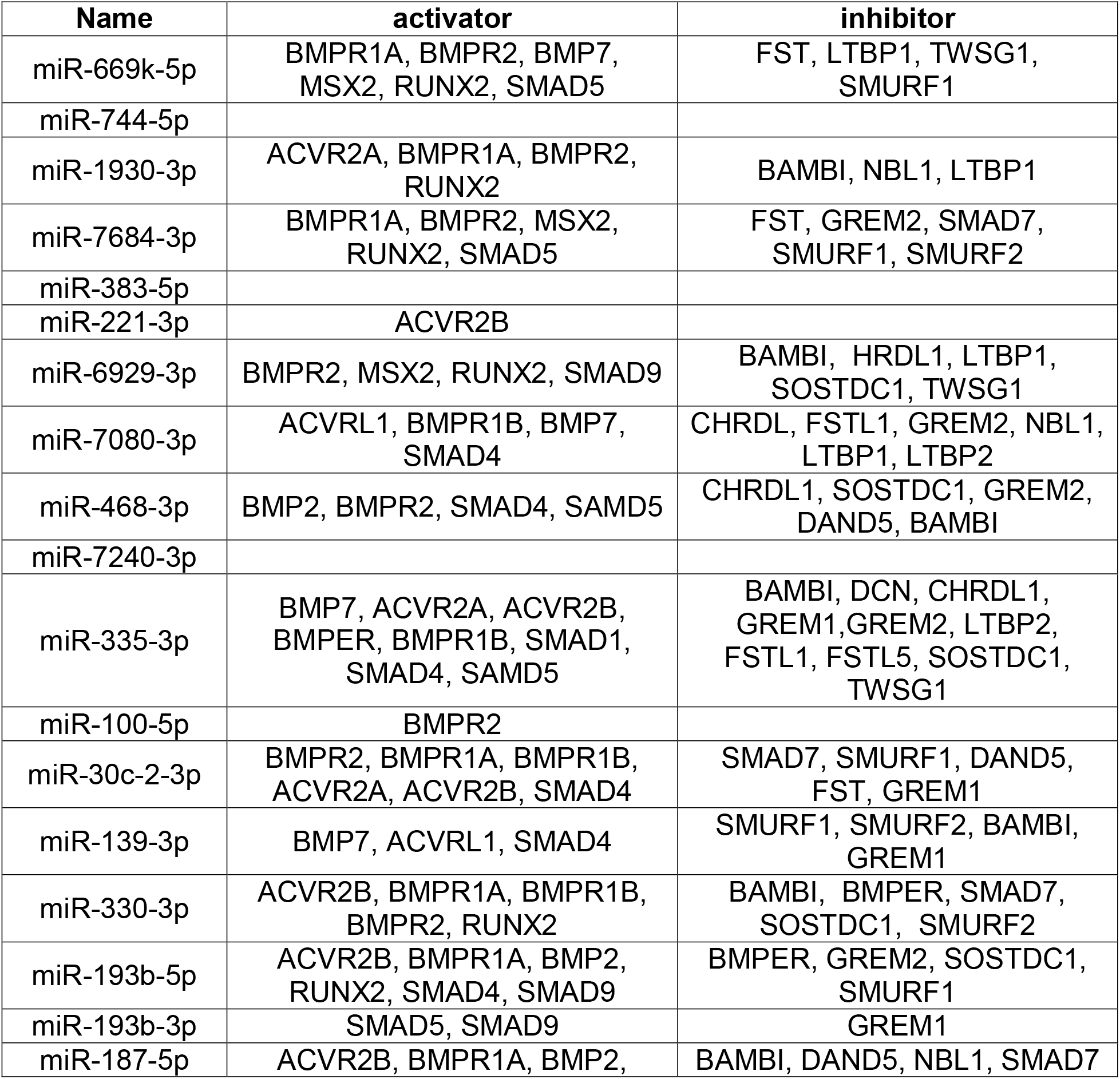
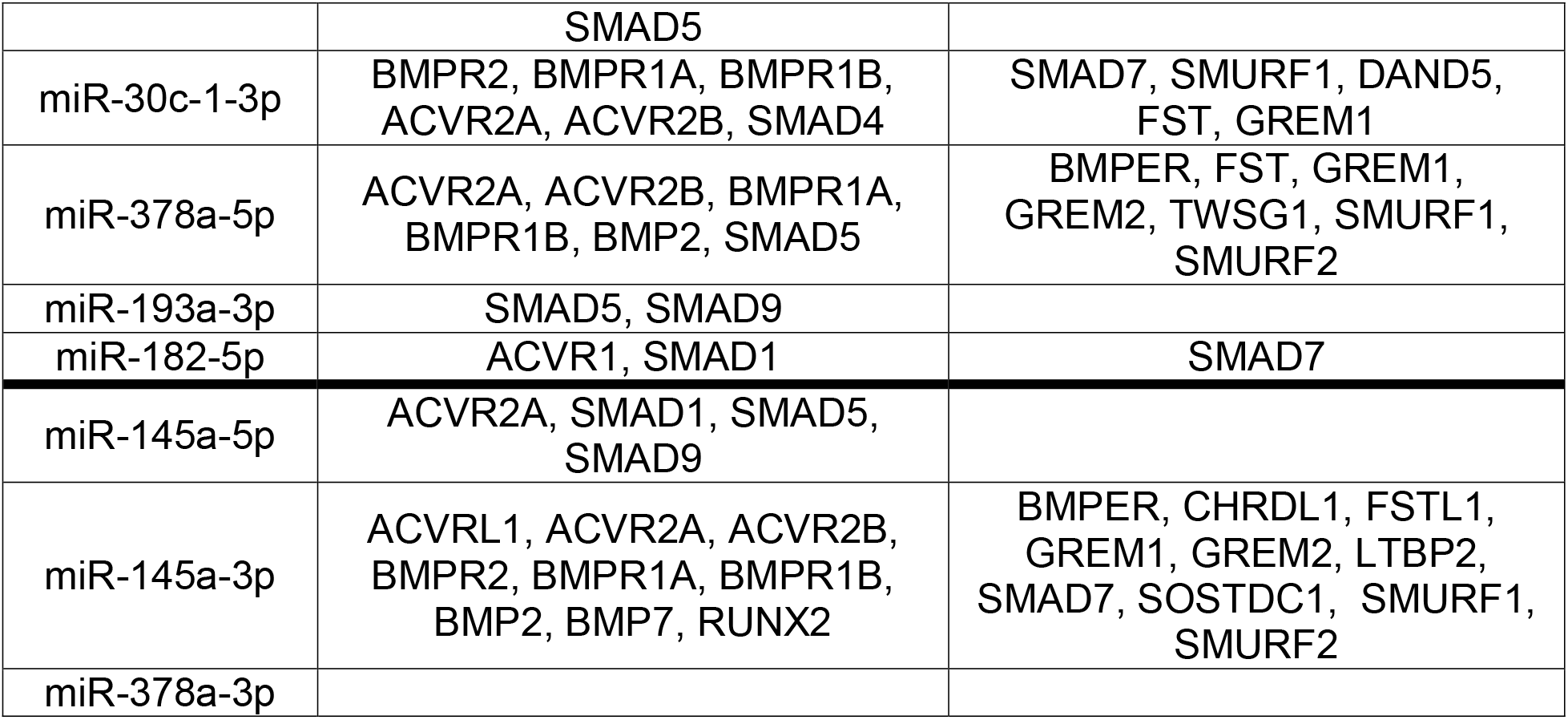
Genes targeted by selected miRNAs that may influence aortic calcification. Twenty-five miRNAs were selected for increased scrutiny because they were differentially regulated to a significant extent in both male and female *Klotho* mutant homozygotes or affected calcium levels in the *in vivo* gain-of-function assay (Table 4, Table S2, GSE135759, Fig. 6 and 7). We list the known BMP signaling modulators (Table S3) or the master osteogenic regulators (MSX2, HGNC:7392 and RUNX2, HGNC:10472) that TargetScan predicted were targeted by each miRNA. Genes have been grouped into primarily activating or inhibiting functional categories. All but four of these differentially regulated miRNAs (miR-744-5p, miR-383-5p, miR-7240-3p, miR-378a-3p) were predicted to target genes that modulate osteogenic differentiation.

### MicroRNA-145 and microRNA-378a attenuation in Klotho mutant mice

We selected two microRNAs to test the principle that a miRNA targeting several members of the BMP signaling pathway would reduce KLOTHO deficiency-associated calcification. Two miRNAs were selected whose abundance was reduced in Klotho mutants. To increase the potential for translational studies, each was predicted to target sequences within the transcripts encoding BMP2 and SMAD proteins that are conserved between mice and humans (Table 6).

**Table 6.**
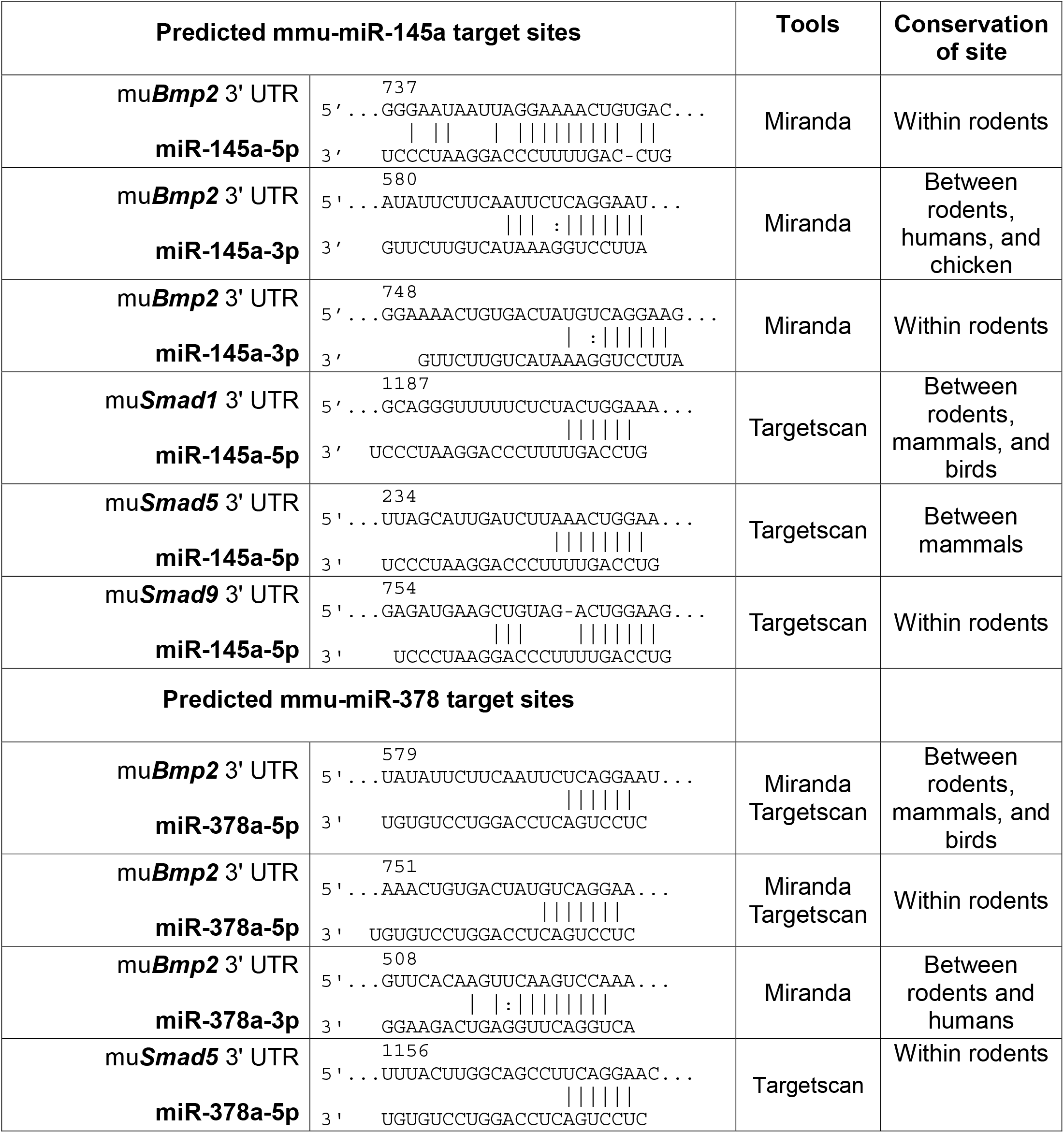
Sequence alignments between miR-145a-5p, miR-145a-3p, miR-378a-5p and miR-378a-3p and mRNAs encoding BMP signaling proteins

The abundance of miR-378a-5p, was sharply decreased in both male and female *Klotho* mutant mice (53% and 38% respectively, Table 4, Table S2). RT-PCR validated our microarray results and confirmed that the abundance of miR-378a-5p and −3p in *Klotho* mutant mice was half that present in healthy control aorta (Fig. 3C, D). TargetScan predicted that miR-378a-5p may target 13 proteins involved in BMP signaling including *Bmp2* and *Smad5* (Table 5). In contrast, TargetScan failed predict that miR-378a-3p would impact BMP signaling (Table 5). Although the abundance of miR-378a-5p did not differ significantly in males and females, a significant difference in miR-378a-3p levels was observed (Fig. 3D). In this report, we focus on miRNAs that may regulate pathological osteogenesis in the *Klotho* model. Because miR-378a-3p is not predicted to strongly interact with BMP signaling genes, we will address this observation in a separate ongoing study of how sex impacts miRNA levels.

**Fig. 3.**
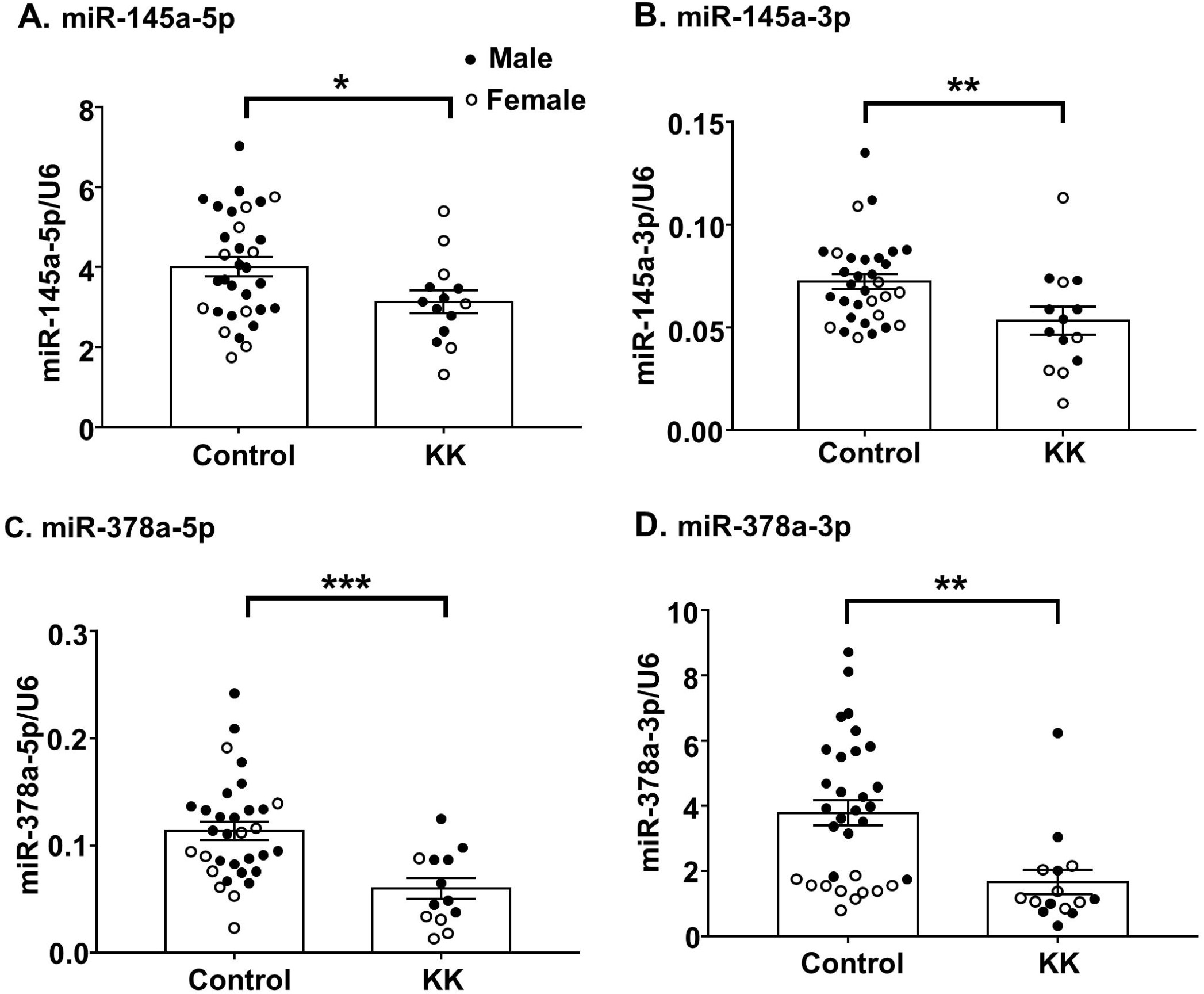
MiR-145 and miR-378a abundances are reduced in the aorta of *Klotho* homozygous mutant mice. Relative miRNA levels in healthy controls (*Kl*^*+/+*^ and *Kl*^*kl*/*+*^) or *Klotho* homozygous (*Kl^kl/kl^*) mice (**KK**) were assessed by RT PCR normalized against the corresponding U6 expression **(A-D)**. The bars represent the mean value +/− SEM and solid and hollow circles represent individual male and female values, respectively. * = *p* < 0.05, ** = *p* < 0.01, *** = *p* < 0.005. The abundances of miR-145-5p and −3p and miR-378a-5p did not differ significantly in males and females. However, miR-378a-3p levels differed significantly in control mice (p < 0.00001), but not in *Klotho* homozygotes.

Our microarray measurements showed that the abundance of miR-145a-5p was reduced by 20% in male *Klotho* homozygous mutant mice as compared to healthy control males (Table S2). RT-PCR indicated that levels of both miR-145a-5p and −3p were significantly reduced in *Klotho* mutant mice of both sexes (Fig. 3A, B). We selected this miRNA for further assessment because, miR-145a is predicted to target *Bmp2*, *Smad1*, *5* and *9* (Table 5, 6, (47)), was also down-regulated in *ApoE* null female mice with partial nephrectomy (45), and improves atherosclerotic symptoms in a mouse model of hyperlipidemia (27). The experiments described below test the impact of restoring miR-145 abundance in an alternative setting of kidney failure with accelerated aging.

### Forced overexpression of mir-145 and mir-378a in Klotho mutant mice

We used a vascular smooth muscle cell (VSMC)-specific *mSm22a* promoter-driven lentivirus to increase the abundance of miR-145 and miR-378a in the aortas of *Klotho* mutant homozygotes (Fig. 4A, (27)). To evaluate the efficacies and persistence of lentiviral transduction, RT-PCR was used to test the aortic levels of miR-145 and miR-378a in mice injected with viruses bearing these miRNAs relative to empty virus vectors. The miR-145 virus raised miR-145a-5p and miR-145a-3p abundances by 1.4±0.8 and 2.0±0.01 fold, respectively, relative to empty virus (Fig. 4B, C). Similarly, miR-378a-5p abundance increased by 1.7±0.04 fold (Fig. 4D). Although unlikely to directly influence BMP signaling, the fact that miR-378a-3p abundance increased significantly in both males and females (Fig. 4E, F) further verifies that overexpression of the precursor RNA occurred in the aorta. Thus this delivery dosage and method successfully augmented aortic miR-145 and miR-378a levels in *Klotho* mutant mice.

**Fig. 4.**
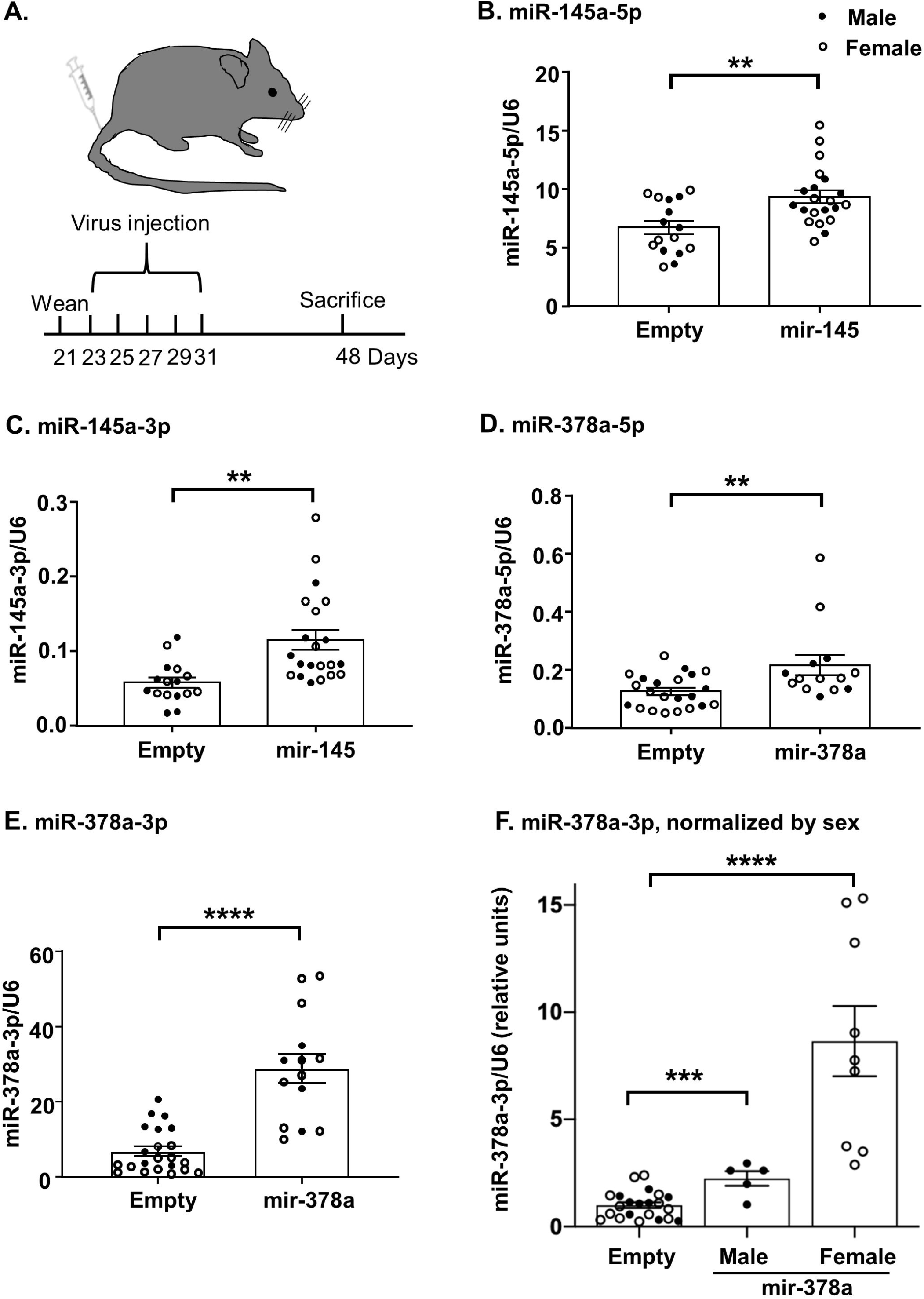
Vascular smooth muscle cell (VSMC)-specific lentivirus treatment increases mir-145 and mir-378a abundance in the aorta of *Klotho* mutant mice. **A.** Weaned *Klotho* mutant homozygous mice were treated with empty virus or virus bearing miR-145 or miR-378a pre-miRNAs. This lentiviral construct was previously shown to deliver miR-145 specifically to aortic VSMCs (27). The miR-145a-5p **(B)**, miR-145a-3p **(C)** and miR-378a-5p **(D)**, miR-378a-3p **(E)** abundances were evaluated by real-time PCR and normalized against the corresponding U6 abundance. The bars represent the mean value +/− SEM. Solid and hollow circles represent individual male and female values, respectively. ** = *p* < 0.01, *** = *p* < 0.005, **** = *p* < 0.001. The induction of miR-145a-5p and −3p and miR-378a-5p levels did not differ significantly between sexes and are presented as values normalized to U6 levels. However, as shown in Fig. 3, miR-378a-3p levels differed in control males and females. Consequently, miR-378a-3p levels are also presented normalized by sex to the values observed in mice exposed to empty virus **(F)**.

### Increased expression of miR-145 and miR-378a reduced BMP signaling and aortic calcification

Four different proteins involved in BMP signaling: *Bmp2, Smad1, 5 and 9* are predicted to be targeted by miR-145 (Table 5, 6). Therefore, we hypothesized that increased miR-145 levels would repress the synthesis of these pro-osteogenesis factors and ameliorate pathological calcification. Having successfully augmented the level of miR-145 with lentivirus delivery (Fig. 4), we evaluated the impact of this treatment on *Bmp2* and *Smad1, 5* and *9* RNA levels with RT-PCR. The *Bmp2* and *Smad5* RNA levels in aortas from mice injected with miR-145-bearing virus fell to 48±0.03% and 65±0.006% (p=0.002 and 0.048) that of control aortas from mice injected with empty virus (Fig. 5A, C). The miR-145 virus did not significantly change the abundances of the *Smad1* (Fig. 5B) and *Smad9* (Fig. 5D).

**Fig. 5.**
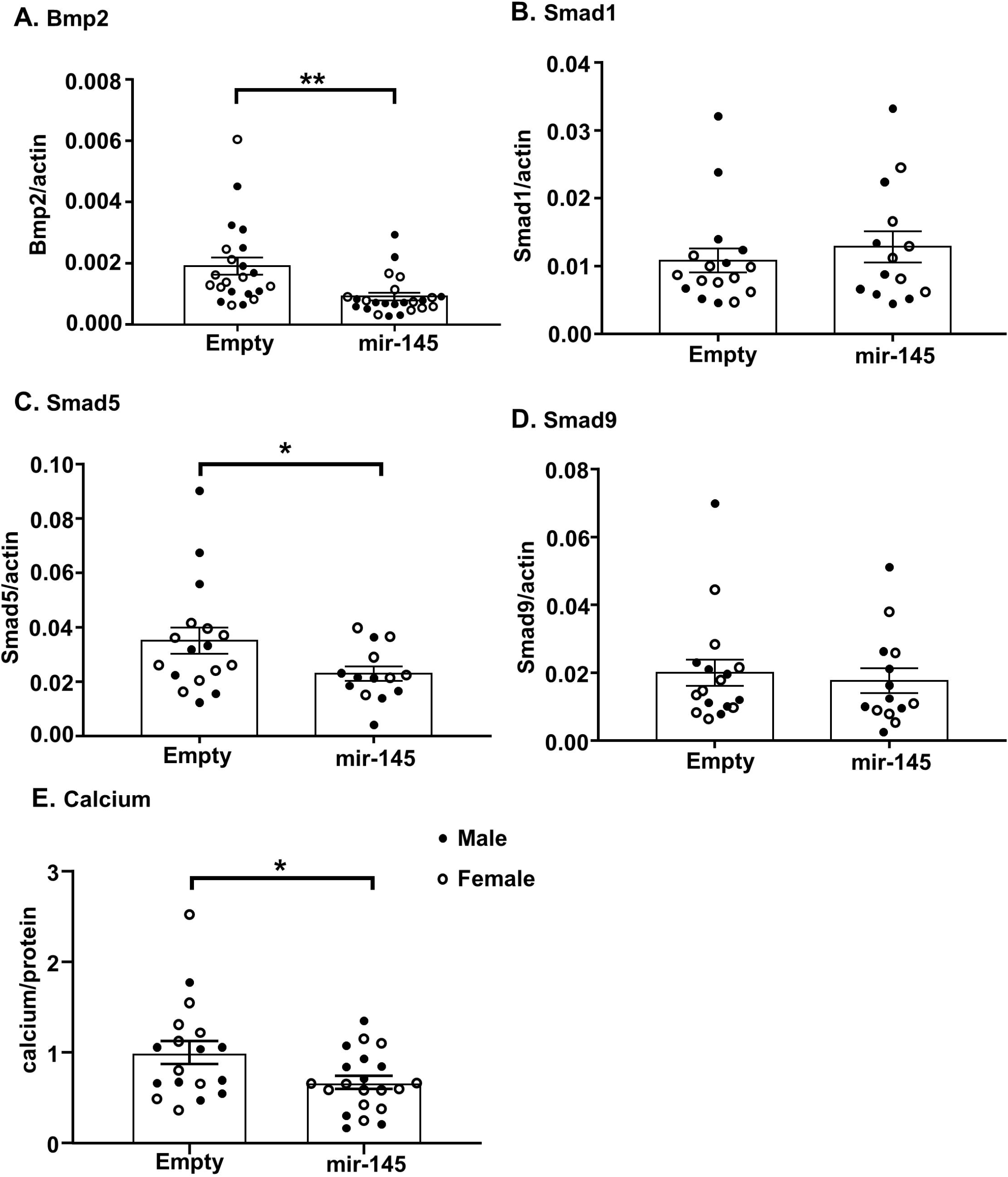
Restoring MiR-145 levels reduces the abundance of RNAs encoding BMP signaling factors and aortic calcification. Increased mir-145 abundance in *Klotho* mutant mice significantly decreased *Bmp2* **(A)** and *Smad5* **(C)** RNA levels and calcium levels **(E)**. *Smad1* **(B)** and *Smad9* **(D)** RNA levels did not change significantly. The bars represent the mean value +/− SEM. Solid and hollow circles represent individual male and female values, respectively. * = *p* < 0.05, ** = *p* < 0.01.

To test if miR-145 overexpression inhibited pathological calcification in *Klotho* homozygous mutant mice, we compared the calcium present in the aortas of *Klotho* homozygous mutant mice injected with empty lentivirus vector or with virus bearing miR-145. The *Klotho* homozygous mutant mice injected with the miR-145 virus had one third less aortic calcium than the empty virus injected mice (67±0.8%, p =0.02, Fig. 5E). Thus, forced miR-145 expression limited pathological calcification in *Klotho* aorta.

The down-regulated miR-378a-5p miRNA is predicted to target *Bmp2* and *Smad5*, but not *Smad1* and *9* (Table 5, 6). Furthermore, published reporter gene studies confirmed that miR-378 directly regulates *Bmp2* (18, 52). The circulating level of miR-378 is reduced in the plasma of patients with coronary heart disease (5) which is consistent with a role in cardiovascular calcification. To evaluate the potiential benefit of restoring miR-378a expression in the aorta of *Klotho* mutant mice, a lentivirus bearing miR-378a was injected. This virus reduced the levels of the target gene RNAs *Bmp2* and *Smad5* to 60±0.02% (p =0.02) and 64±0.4% (p =0.04) respectively of the RNA levels observed in the mice injected with empty virus (Fig. 6A and B). Finally, treatment with the mir-378a-bearing virus reduced aortic calcium levels by a third (63 ±0.6%, p=0.02) relative to levels in mice exposed to the empty virus (Fig. 6C). Thus as observed for miR-145, forced expression of miR-378a significantly ameliorated pathological calcification of the aorta.

**Fig. 6.**
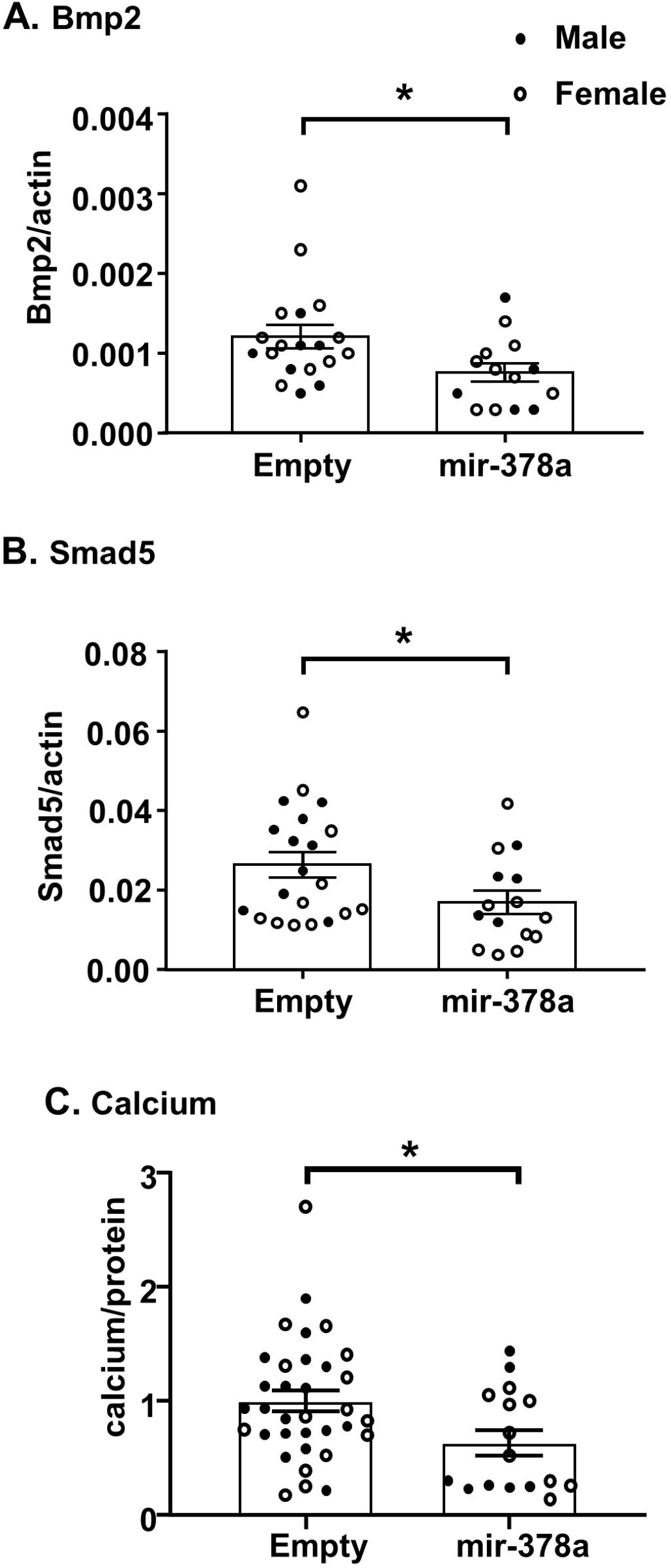
Restoring MiR-378a levels reduces the abundance of RNAs encoding BMP signaling factors and aortic calcification. Increased mir-378a abundance in *Klotho* mutant mice significantly decreased *Bmp2* **(A)**, *Smad5* **(B)** RNA and calcium levels **(C)**. The bars represent the mean value +/− SEM. Solid and hollow circles represent individual male and female values, respectively. * = *p* < 0.05.

## Discussion

Large datasets comparing organs with different clinically relevant physiologies are important starting points for empirical investigations. Here, we provide comprehensive profiles of differentially regulated miRNAs in healthy *vs*. calcified aorta. Our goal is to improve our understanding of regulatory differences leading to vascular calcification. We contribute miRNA profiles in aortas from control healthy mice and in *Klotho* homozygous mutant mice with renal disease to the public databases. KLOTHO deficiency leads to renal impairment and pathological osteogenesis within the vasculature. This vascular bone formation involves pathologically increased BMP signaling. Consequently, we initially focused on the subset of differentially regulated miRNAs that impact BMP signaling and key osteogenesis regulators. Most importantly, we demonstrated that selected miRNAs can modulate the BMP2 ligand and proteins involved in BMP signaling and calcification *in vivo*. An additional strength of our study is the rigorous inclusion of samples from both male and female mice in all experiments.

The abundances of many miRNAs were changed in the same direction in both male and female mice (Table 4). However, the statistical significance of the change for some miRNAs failed to reach our cutoff of p<0.05 in both sexes. Consequently, only a small fraction of miRNAs whose levels changed to a statistically significant level were regulated similarly in both male and female mice (Fig. 2C). This quantitative difference is partly explained by a limitation of our study whereby fewer female *Klotho* homozygotes were available (Table 1).

However, we also observed qualitative differences in the miRNA profiles of males and females. For example, the abundances of miR-3069-3p, miR-7019-5p, miR-7685-3p, miR-679-5p, miR-370-5p, miR-7665-3p, miR-466m-3p, miR-17-5p and miR-181c-3p changed in opposite directions in males and females (Table 4). Interestingly, a class of miRNAs that modulate epithelial mesenchymal transition (40) were highly up-regulated exclusively in the aorta of female mice. MiR-205-5p and −3p and five members of the miR-200 family (miR-141-5p and −3p, miR-200a-5p and −3p, miR-429-3p) were up-regulated by as much as 65-fold in *Klotho* mutant females but were unchanged in males (Table S2). Epithelial mesenchymal transition influences vascular calcification (9, 41). Both quantitative and qualitative differences in aortic miRNA profiles in each sex may be highly clinically relevant because there are significant disparities in incidence, prognosis, and response to treatments for arterial diseases between men and women (13). We are separately investigating the sex-associated differences in the context of a study aimed at clarifying the impact of sex on BMP signaling (42).

We selected two miRNAs, miR-145 and miR-378, to test the principle that miRNAs targeting members of the BMP signaling pathway would reduce aortic calcification. We used the *Klotho* hypomorphic model because the swift pace of aortic calcification in the Klotho mutant homozygotes facilitates testing therapeutic approaches to reducing calcification. In contrast, other models are experimentally time-consuming. For example, mice with genetically sensitized hyperlipidemia backgrounds must be fed special diets for months (29, 35). In *Klotho* mutant mice with hyperphosphatemia, elevated aortic BMP signaling and calcification is observed at 6-7 weeks of age (Fig. 1, (17)). In this model, only 10 days of exposure to viruses overexpressing miR-145 and miR-378 limited aortic calcification (Fig. 5, 6). Studies using *Klotho* mice with renal disease also are relevant to human biology, because KLOTHO deficiency is associated with chronic kidney disease that promotes valve and vascular calcification in people (2, 3, 20, 23). This work underscores the great potential for miRNA-based therapies in treating calcification pathologies.

Restoration of aortic miR-145 levels has also been tested in a different model of vascular calcification. Forced expression of miR-145 reduced atherosclerotic plaque size, increased plaque stability, and promoted a contractile cell phenotype in the *ApoE*, high fat diet mouse model of hyperlipidemia (27). Our demonstration of reduced calcification in *Klotho* mutant homozygous mice with forced miR-145 (Fig. 5) indicates that miR-145 also attenuates vascular disease in a model of atherosclerosis caused by hyperphosphatemia. The effectiveness of this microRNA in these highly dissimilar physiological situations suggest that miR-145 directly promotes vascular health and inhibits pathological osteogenesis.

Like miR-145, miR-378a was significantly down-regulated in blood samples from patients with coronary artery disease compared to healthy subjects (48). Furthermore, both miRNAs target regions of the BMP2 and other BMP signaling genes that are highly conserved between mice and humans (Table 6, (18, 52)). Interestingly, both miRNAs are subject to editing whereby specific adenosines are post-transcriptionally converted to inosine (44, 55). Inosine base pairs cytosine, not thymine. Manual sequence inspection indicates that editing would increase the complementarity of miR-145 and miR-378a for sites within both the mouse and human *BMP2* messages. Although editing may significantly change miRNA/message interactions, current databases and prediction tools do not facilitate global predictions of how edited miRNAs may impact signaling pathways.

In summary, we have provided a database of the miRNAs that are differentially expressed in healthy mouse aorta relative to aorta calcified due to defective mineral metabolism and kidney function. These databases and all experiments described in this report include samples from both male and female mice. We also demonstrated that forced expression of miR-145 and miR-378 inhibited RNAs encoding BMP signaling proteins and chronic kidney disease-associated aortic calcification. MiRNA-145 and miR-378 are candidates for translational studies investigating therapies to block vascular calcification.

## Acknowledgements

We warmly thank Youhua Zhu for her diligent technical assistance, Dr. Diane Garsetti and Yue Wang for critically reading the manuscript, and Lindsey Hernandez for bioinformatics assistance. We appreciate the technical assistance provided by the NJMS Genomics Center.

Tapan A. Shah’s present address is Advanced Cell Diagnostics, 7707 Gateway Blvd #200, Newark, CA 94560

Funding was provided by grants from the National Heart, Lung, and Blood Institute (R01HL114751) and National Institutes of Aging (R56AG050762) and from the NJMS Research Core Facilities Matching Funds Small Grants Program to MBR and an American Heart Association fellowship (20POST35210235) to YT.

