## supporting file for "MicroRNA Profiles in Calcified and Healthy Aorta Differ: Therapeutic Impact of miR-145 and miR-378"

**Running title:** *miRNA replacement in aortic calcification*

Ying Tang^1, †^, Tapan A. Shah^1, †, ‡^, Edward J. Yurkow^2^, and Melissa B. Rogers^1, ‡^

^1^Rutgers - New Jersey Medical School, Microbiology, Biochemistry, & Molecular Genetics, Newark, NJ

^2^Rutgers University Molecular Imaging Center (RUMIC), Rutgers University, Piscataway, NJ

^†^Co-first authors

^‡^Present Address: Advanced Cell Diagnostics, 7707 Gateway Blvd #200, Newark, CA 94560

^§^To whom should correspondence and reprint request be addressed to: Melissa B. Rogers, Ph.D., Microbiology, Biochemistry & Molecular Genetics, Rutgers - NJ Medical School (NJMS), Center for Cell Signaling, Room F1216, 205 South Orange Ave., Newark, NJ 07103., telephone: 973 972 2984

Fig. S1. Experimental Controls.

Table S1. *Klotho* wild type and *Klotho* heterozygous mice are equivalent controls.

Table S2. Average miRNA abundance in aorta from *Klotho* mutant homozygotes relative to healthy control (p<0.05).

Table S3. Known modulators of BMP signaling.

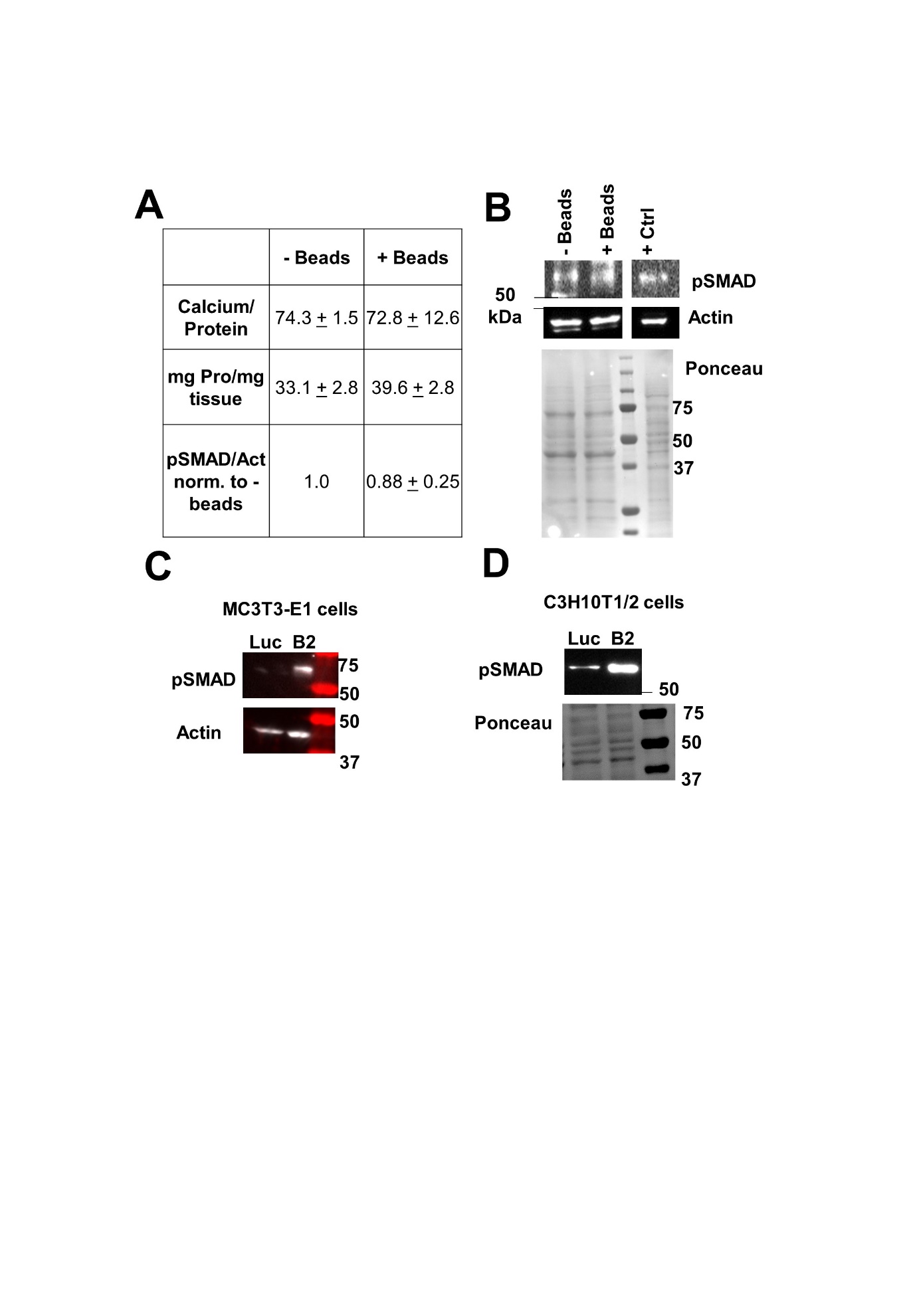

**Fig. S1. Experimental Controls.** **A, B.** Grinding tissue with glass beads does not significantly affect calcium or protein yield. Whole aorta was ground in liquid nitrogen either with or without glass beads. Frozen ground powder tissue was split two ways for protein and calcium assays. **A.** Effect of glass beads on calcium, protein and pSMAD1/5/9 levels. Duplicate measurements are presented with range. Experiments were repeated twice with similar results. **B.** Representative blots showing pSMAD1/5/9, actin levels and the Ponceau S stained membrane after transfer. The positive control lane (+ Ctrl) was loaded with lysate from MC3T3-E1 cells transfected with a *Bmp2* expression plasmid. **C, D.** Validation of the phospho-SMAD1/5/9 antibody. MC3T3-E1 (**C**) and C3H10T1/2 (**D**) cells were transfected with an expression plasmid encoding BMP2 (B2) or luciferase (Luc) (Kruithof *et al.*, 2011). Cells were then lysed in RIPA buffer and subjected to western blotting as described in the experimental procedures section. These representative blots show that pSMAD1/5/9 levels were induced in cells transfected with the *Bmp2* expressing plasmid relative to the luciferase plasmid. Actin levels (**C**) and a Ponceau S (**D**) stained membrane are shown as loading controls.

**Table S1. *Klotho* wild type and *Klotho* heterozygous mice are equivalent controls.** Wildtype *Bmp2^+/+^* control (Wt), *Bmp2Kl/+* (K+), Average (Ave.), Standard deviation (SD), number of mice measured (n)

|  | **MALES** | | | | | |  | **FEMALES** | | | | | |
| --- | --- | --- | --- | --- | --- | --- | --- | --- | --- | --- | --- | --- | --- |
| **Klotho genotype** | **Wt** | | | **K+** | | |  | **Wt** | | | **K+** | | |
| **Two-way Anova**  ***p* value** | **0.29** | | | | | |  | **0.21** | | | | | |
| **Parameters** | **Ave.** | **SD** | **n** | **Ave.** | **SD** | **n** |  | **Ave.** | **SD** | **n** | **Ave.** | **SD** | **n** |
| *Bmp2 RNA* | 0.72 | 0.24 | 7 | 0.78 | 0.46 | 16 |  | 0.57 | 0.09 | 5 | 0.89 | 0.31 | 6 |
| pSMAD/Total SMAD | 0.61 | 0.30 | 4 | 0.58 | 0.05 | 2 |  | 1.16 | 0.50 | 3 | 1.15 | 0.79 | 4 |
| Calcium/Protein | 42.1 | 21.8 | 7 | 51.2 | 21.2 | 11 |  | 64.8 | 23.4 | 5 | 53.2 | 14.5 | 3 |
| Body Weight (g) | 22.3 | 2.8 | 13 | 22.3 | 1.4 | 18 |  | 18.2 | 1.6 | 8 | 18.8 | 2.3 | 10 |
| Heart Weight (g) | 0.097 | 0.027 | 13 | 0.111 | 0.021 | 16 |  | 0.082 | 0.014 | 8 | 0.093 | 0.013 | 10 |
| Heart / Body Weight | 0.0044 | 0.0012 | 13 | 0.0049 | 0.0010 | 15 |  | 0.0045 | 0.0004 | 8 | 0.0049 | 0.0006 | 10 |
| Age (d) at euthanasia | 47.8 | 3.7 | 13 | 48.9 | 1.8 | 19 |  | 49.4 | 3.1 | 8 | 48.5 | 3.5 | 10 |

**Table S2. Average miRNA abundance in aorta from *Klotho* mutant homozygotes relative to healthy control (p<0.05).** Raw data is available in the Gene Expression Omnibus (<https://www.ncbi.nlm.nih.gov/geo/>, GSE135759).

| Male | % control | Female | % control |
| --- | --- | --- | --- |
| miR-16-1-3p | 3852 | miR-205-5p | 6508 |
| miR-466j | 2293 | miR-6359 | 1625 |
| miR-466m-5p | 1906 | miR-205-3p | 1121 |
| miR-669m-5p | 1906 | miR-1947-3p | 1013 |
| miR-466f | 1630 | miR-675-3p | 977 |
| miR-711 | 1626 | miR-291b-5p | 800 |
| miR-669b-5p | 1551 | miR-141-3p | 780 |
| miR-466h-5p | 1496 | let-7f-1-3p | 680 |
| miR-7030-5p | 1370 | miR-429-3p | 617 |
| miR-195a-3p | 1210 | miR-489-3p | 561 |
| miR-297a-5p | 1102 | miR-200c-3p | 560 |
| miR-466f-5p | 1100 | miR-200a-5p | 480 |
| miR-669f-5p | 982 | miR-539-5p | 465 |
| miR-669l-5p | 966 | miR-3092-3p | 454 |
| miR-7005-5p | 931 | miR-1930-3p | 448 |
| miR-7033-5p | 907 | miR-1946b | 423 |
| miR-669e-5p | 893 | miR-381-5p | 421 |
| miR-696 | 679 | miR-296-3p | 416 |
| miR-467h | 648 | miR-6241 | 406 |
| miR-669o-5p | 646 | miR-7047-3p | 396 |
| miR-5620-5p | 643 | miR-669k-5p | 395 |
| miR-7221-3p | 631 | miR-7049-5p | 383 |
| miR-3082-5p | 631 | miR-7079-5p | 377 |
| miR-669d-5p | 630 | miR-183-3p | 357 |
| miR-669k-5p | 624 | miR-7004-5p | 351 |
| miR-665-5p | 620 | miR-6546-3p | 348 |
| miR-5620-3p | 610 | miR-7241-5p | 339 |
| miR-346-3p | 552 | miR-376a-3p | 338 |
| miR-5100 | 532 | miR-181c-3p | 320 |
| miR-7686-5p | 529 | let-7b-3p | 312 |
| miR-7648-3p | 502 | miR-191-3p | 307 |
| miR-6991-5p | 455 | miR-1946a | 307 |
| miR-714 | 452 | miR-744-5p | 304 |
| miR-762 | 434 | miR-203-5p | 301 |
| miR-466c-5p | 434 | miR-19a-3p | 285 |
| miR-15a-3p | 416 | miR-18a-3p | 277 |
| miR-6968-5p | 392 | miR-5129-3p | 263 |
| miR-7235-5p | 392 | miR-141-5p | 250 |
| miR-122-5p | 370 | miR-7682-3p | 248 |
| miR-7023-5p | 360 | miR-7684-5p | 248 |
| miR-1224-5p | 347 | miR-6958-5p | 248 |
| miR-1894-3p | 342 | miR-8103 | 246 |
| miR-6349 | 335 | miR-7684-3p | 238 |
| miR-21a-5p | 330 | miR-7082-3p | 236 |
| miR-574-5p | 326 | miR-7077-5p | 229 |
| miR-7654-3p | 319 | miR-7669-3p | 223 |
| miR-5130 | 314 | miR-702-3p | 220 |
| miR-8119 | 310 | miR-290b-5p | 204 |
| miR-6909-5p | 299 | miR-106a-5p | 195 |
| miR-6980-5p | 296 | miR-704 | 194 |
| miR-7016-5p | 284 | miR-17-5p | 194 |
| miR-6912-5p | 282 | miR-468-3p | 190 |
| miR-146b-5p | 282 | miR-673-3p | 190 |
| miR-7011-5p | 279 | miR-200c-5p | 188 |
| miR-2137 | 272 | miR-7057-3p | 185 |
| miR-92b-5p | 266 | miR-3544-3p | 182 |
| miR-6921-5p | 265 | miR-146b-3p | 181 |
| miR-3547-5p | 259 | miR-18b-5p | 181 |
| miR-6971-5p | 257 | let-7i-3p | 178 |
| miR-669a-5p | 256 | miR-383-5p | 176 |
| miR-669p-5p | 256 | miR-7008-5p | 171 |
| miR-6240 | 248 | miR-92a-1-5p | 170 |
| miR-7069-5p | 246 | miR-6929-3p | 170 |
| miR-5119 | 233 | miR-465a-5p | 167 |
| miR-744-5p | 232 | miR-147-3p | 167 |
| miR-8101 | 227 | miR-292-3p | 162 |
| miR-1982-5p | 224 | miR-7074-3p | 161 |
| miR-1892 | 218 | miR-3098-5p | 161 |
| miR-7045-5p | 217 | miR-450b-5p | 159 |
| miR-7083-5p | 216 | miR-1983 | 158 |
| miR-7118-5p | 215 | miR-7007-3p | 153 |
| miR-214-3p | 214 | miR-297c-5p | 150 |
| miR-5128 | 213 | miR-3109-5p | 149 |
| miR-1930-3p | 210 | miR-7115-3p | 149 |
| miR-1249-5p | 207 | miR-221-3p | 148 |
| miR-7036-5p | 206 | miR-6959-5p | 148 |
| miR-7684-3p | 202 | miR-712-3p | 148 |
| miR-7075-5p | 196 | miR-7087-3p | 147 |
| miR-7070-5p | 192 | miR-7080-3p | 146 |
| miR-3069-3p | 190 | miR-7227-3p | 144 |
| miR-7048-5p | 184 | miR-1930-5p | 144 |
| miR-6391 | 183 | miR-7054-5p | 143 |
| miR-5132-5p | 178 | miR-547-5p | 143 |
| miR-3470a | 177 | miR-7213-5p | 141 |
| miR-7085-5p | 177 | miR-743a-5p | 139 |
| miR-6935-5p | 176 | let-7f-2-3p | 139 |
| miR-328-5p | 172 | miR-7225-3p | 138 |
| miR-7036b-3p | 169 | miR-7092-3p | 138 |
| miR-7052-5p | 168 | miR-707 | 138 |
| miR-6394 | 167 | miR-6965-3p | 138 |
| miR-6769b-5p | 167 | miR-467g | 137 |
| miR-382-5p | 163 | miR-5103 | 137 |
| miR-7034-5p | 163 | miR-7651-3p | 136 |
| miR-6988-5p | 160 | miR-217-5p | 135 |
| miR-7042-5p | 158 | miR-293-5p | 134 |
| miR-383-5p | 153 | miR-135b-3p | 132 |
| miR-185-5p | 152 | miR-6996-3p | 131 |
| miR-6386 | 149 | miR-6976-5p | 130 |
| miR-680 | 147 | miR-7687-3p | 129 |
| miR-6956-5p | 146 | miR-804 | 128 |
| miR-3473g | 146 | miR-488-5p | 128 |
| miR-7019-5p | 146 | miR-21b | 128 |
| miR-760-3p | 146 | miR-33-5p | 127 |
| miR-221-3p | 144 | miR-6920-3p | 126 |
| miR-7010-3p | 144 | miR-7678-5p | 125 |
| miR-6979-5p | 144 | miR-6937-3p | 125 |
| miR-6418-5p | 142 | miR-7008-3p | 125 |
| miR-6929-3p | 142 | miR-466m-3p | 124 |
| miR-3535 | 141 | miR-7665-3p | 124 |
| miR-6516-3p | 139 | miR-505-3p | 123 |
| miR-698-5p | 138 | miR-124-5p | 123 |
| miR-7676-3p | 134 | miR-6387 | 123 |
| miR-6348 | 134 | miR-8092 | 123 |
| miR-7080-3p | 133 | miR-6992-3p | 121 |
| miR-1195 | 131 | miR-6377 | 121 |
| let-7b-5p | 130 | miR-7029-5p | 118 |
| miR-5617-5p | 130 | miR-6404 | 118 |
| miR-7668-3p | 129 | miR-592-3p | 116 |
| miR-468-3p | 126 | miR-7240-3p | 115 |
| miR-218-1-3p | 126 | miR-24-3p | 112 |
| miR-6999-3p | 125 | miR-6906-3p | 112 |
| miR-6966-5p | 124 | miR-6398 | 87 |
| miR-7119-5p | 123 | miR-7650-5p | 86 |
| miR-466h-3p | 123 | miR-3094-5p | 85 |
| miR-3084-5p | 123 | miR-135a-5p | 84 |
| miR-7685-3p | 122 | miR-6951-5p | 84 |
| miR-6983-3p | 122 | miR-6961-5p | 83 |
| miR-679-5p | 122 | miR-7222-5p | 83 |
| miR-3961 | 121 | miR-6361 | 82 |
| let-7c-5p | 120 | miR-7685-3p | 82 |
| miR-107-5p | 119 | miR-216c-5p | 82 |
| miR-7073-3p | 119 | miR-679-5p | 82 |
| miR-3963 | 118 | miR-7678-3p | 81 |
| miR-3103-5p | 118 | miR-370-5p | 81 |
| miR-144-3p | 118 | miR-6415 | 81 |
| miR-6935-3p | 118 | miR-758-3p | 81 |
| miR-466b-5p | 117 | miR-551b-3p | 80 |
| miR-466o-5p | 117 | miR-653-5p | 80 |
| miR-370-5p | 117 | miR-880-5p | 80 |
| miR-374c-3p | 116 | miR-1264-3p | 80 |
| miR-6416-3p | 115 | miR-669e-3p | 80 |
| miR-219c-3p | 113 | miR-7675-3p | 79 |
| miR-6411 | 112 | miR-7061-3p | 79 |
| miR-7240-3p | 110 | miR-7232-5p | 78 |
| miR-7064-3p | 110 | miR-1938 | 77 |
| miR-3964 | 92 | miR-7656-3p | 77 |
| miR-3097-3p | 89 | miR-8108 | 77 |
| miR-3075-3p | 89 | miR-3074-5p | 77 |
| miR-7054-3p | 88 | miR-7237-5p | 76 |
| let-7a-2-3p | 88 | miR-7029-3p | 76 |
| miR-7065-3p | 87 | miR-344c-5p | 76 |
| miR-1258-5p | 87 | miR-7044-3p | 76 |
| miR-6922-3p | 87 | miR-3065-3p | 75 |
| miR-325-3p | 87 | miR-302c-5p | 74 |
| miR-6940-5p | 86 | miR-7685-5p | 74 |
| miR-7665-3p | 85 | miR-1969 | 73 |
| miR-202-3p | 85 | miR-99b-5p | 73 |
| miR-539-3p | 85 | miR-758-5p | 73 |
| miR-341-5p | 85 | miR-1198-5p | 73 |
| miR-717 | 84 | miR-335-3p | 72 |
| miR-143-3p | 84 | miR-883a-3p | 72 |
| miR-466m-3p | 84 | miR-7649-3p | 71 |
| miR-335-3p | 84 | miR-30a-5p | 69 |
| miR-3057-3p | 84 | miR-361-5p | 65 |
| miR-670-3p | 83 | miR-466b-3p | 65 |
| miR-3086-3p | 83 | miR-466c-3p | 65 |
| miR-23b-3p | 83 | miR-466p-3p | 65 |
| miR-7657-5p | 82 | miR-320-3p | 64 |
| miR-145a-5p | 82 | miR-152-3p | 63 |
| miR-5710 | 81 | miR-2136 | 63 |
| miR-1247-5p | 81 | miR-100-5p | 62 |
| miR-107-3p | 81 | miR-30d-5p | 61 |
| miR-3101-5p | 80 | miR-351-5p | 60 |
| miR-99a-5p | 80 | miR-7660-3p | 59 |
| miR-7091-5p | 80 | miR-483-5p | 58 |
| miR-7015-3p | 80 | miR-6994-5p | 58 |
| miR-8114 | 79 | miR-126a-3p | 57 |
| miR-19a-5p | 78 | miR-7051-5p | 57 |
| miR-3074-2-3p | 78 | miR-7019-5p | 57 |
| miR-34a-3p | 77 | miR-28a-3p | 56 |
| miR-425-5p | 77 | miR-195a-5p | 54 |
| miR-377-3p | 76 | miR-30c-2-3p | 54 |
| miR-17-5p | 76 | miR-378a-3p | 53 |
| miR-100-5p | 75 | miR-187-3p | 50 |
| miR-16-5p | 74 | miR-182-5p | 50 |
| miR-15a-5p | 72 | miR-30a-3p | 50 |
| miR-5624-3p | 72 | miR-151-5p | 50 |
| miR-497-5p | 72 | miR-676-3p | 49 |
| miR-6417 | 71 | miR-3069-3p | 49 |
| miR-300-3p | 69 | miR-151-3p | 48 |
| miR-466n-3p | 68 | miR-877-5p | 47 |
| miR-532-3p | 64 | miR-139-5p | 45 |
| miR-30c-2-3p | 64 | miR-378d | 43 |
| miR-30c-5p | 62 | miR-139-3p | 42 |
| miR-139-3p | 62 | miR-455-3p | 41 |
| miR-345-5p | 61 | miR-378c | 41 |
| miR-199b-5p | 60 | miR-378b | 40 |
| miR-331-3p | 60 | miR-7001-5p | 39 |
| miR-7092-5p | 59 | miR-330-3p | 39 |
| miR-467a-5p | 59 | miR-378a-5p | 38 |
| miR-181d-5p | 57 | miR-150-5p | 36 |
| miR-24-1-5p | 57 | miR-365-1-5p | 36 |
| miR-350-3p | 57 | miR-7662-3p | 35 |
| miR-148a-3p | 57 | miR-3076-5p | 34 |
| miR-194-5p | 57 | miR-193a-5p | 33 |
| miR-330-3p | 56 | miR-193a-3p | 30 |
| miR-574-3p | 55 | miR-1839-3p | 29 |
| let-7e-3p | 55 | miR-193b-3p | 28 |
| miR-708-5p | 54 | miR-30c-1-3p | 28 |
| miR-22-5p | 54 | miR-124-3p | 23 |
| miR-1247-3p | 54 | miR-187-5p | 21 |
| miR-193b-5p | 54 | miR-193b-5p | 16 |
| miR-193b-3p | 54 | miR-1941-5p | 13 |
| miR-187-5p | 53 |  |  |
| miR-30c-1-3p | 53 |  |  |
| miR-378a-5p | 53 |  |  |
| miR-324-3p | 53 |  |  |
| miR-106b-5p | 53 |  |  |
| miR-30e-5p | 52 |  |  |
| miR-411-5p | 52 |  |  |
| miR-345-3p | 52 |  |  |
| miR-339-3p | 52 |  |  |
| miR-409-5p | 52 |  |  |
| miR-154-5p | 52 |  |  |
| miR-423-3p | 51 |  |  |
| miR-1843a-5p | 50 |  |  |
| miR-3065-5p | 50 |  |  |
| miR-339-5p | 49 |  |  |
| miR-193a-3p | 49 |  |  |
| miR-322-3p | 46 |  |  |
| miR-328-3p | 46 |  |  |
| miR-503-5p | 46 |  |  |
| miR-31-3p | 46 |  |  |
| miR-181c-3p | 45 |  |  |
| miR-301a-3p | 45 |  |  |
| miR-5121 | 45 |  |  |
| miR-101b-3p | 45 |  |  |
| miR-3102-3p | 44 |  |  |
| miR-484 | 44 |  |  |
| miR-182-5p | 44 |  |  |
| miR-6239 | 41 |  |  |
| miR-326-3p | 40 |  |  |
| miR-181c-5p | 40 |  |  |
| miR-1940 | 39 |  |  |
| miR-299a-3p | 35 |  |  |
| miR-3068-5p | 34 |  |  |
| miR-1843b-5p | 30 |  |  |
| miR-93-3p | 29 |  |  |
| miR-125b-2-3p | 29 |  |  |
| miR-425-3p | 29 |  |  |
| miR-615-3p | 15 |  |  |

**Table S3. Modulators of BMP signaling** (Faraahi *et al.*, 2019; Geng *et al.*, 2011; Katagiri & Watabe, 2016; Lorda-Diez *et al.*, 2013; Lowery & Rosen, 2018; Shah & Rogers, 2018; Suri *et al.*, 2018). The official gene name and HGNC ID (<https://www.genenames.org/>) are shown. Alternative gene names are in parentheses.

| **Gene symbol** | **HGNC ID** |
| --- | --- |
| **Ligands** | |
| BMP2 | 1069 |
| BMP4 | 1071 |
| BMP7 (OP-1) | 1074 |
| **Receptors** | |
| ACVR1 (SKP1; ALK2; ACVR1A) | 171 |
| ACVR2A (ACTRII) | 173 |
| ACVR2B (ActR-IIB) | 174 |
| ACVRL1 (HHT2; ALK1; HHT) | 175 |
| BMPR2 (BRK-3; T-ALK; BMPR3; BMPR-II) | 1078 |
| BMPR1A (ALK3; CD292) | 1076 |
| BMPR1B (ALK6; CDw293) | 1077 |
| **Intracellular Signal Transducers** | |
| SMAD1 (MADR1; JV4-1) | 6767 |
| SMAD4 (DPC4) | 6770 |
| SMAD5 (Dwfc; JV5-1) | 6771 |
| SMAD9 (SMAD8; SMAD8/9) | 6774 |
| **Intracellular Inhibitors** | |
| SMAD6 (HsT17432) | 6772 |
| SMAD7 | 6773 |
| SMURF1 (KIAA1625) | 16807 |
| SMURF2 | 16809 |
| **Extracellular Inhibitors** | |
| BAMBI (NMA) | 30251 |
| BMPER (Cv2; CRIM3) | 24154 |
| CHRD | 1949 |
| CHRDL1 (NPLN1; CHL) | 29861 |
| DAND5 (FLJ38607; CKTSF1B3; DANTE; GREM3; CER2; DTE; Coco) | 26780 |
| DCN (DSPG2; SLRR1B) | 2705 |
| FST | 3971 |
| FSTL1 (FRP; FSL1; OCC1; OCC-1; tsc36) | 3972 |
| FSTL5 (DKFZp566D234; KIAA1263) | 21386 |
| GREM1(DRM; gremlin; DAND2; HMPS) | 2001 |
| GREM2 (Prdc; FLJ21195; CKTSF1B2; DAND3) | 17655 |
| LTBP1 | 6714 |
| LTBP2 (LTBP3, C14orf141) | 6715 |
| NBL1 (D1S1733E; NB; DAN; NO3; DAND1) | 7650 |
| NOG | 7866 |
| SOST (VBCH; DAND6) | 13771 |
| SOSTDC1 (DKFZp564D206; USAG1; DAND7) | 21748 |
| TWSG1 (TSG) | 12429 |
